# The scheduling of adolescence with Netrin-1 and UNC5C

**DOI:** 10.1101/2023.01.19.521267

**Authors:** Daniel Hoops, Robert F. Kyne, Samer Salameh, Del MacGowan, Radu G. Avramescu, Elise Ewing, Alina T. He, Taylor Orsini, Anais Durand, Christina Popescu, Janet M. Zhao, Kelcie C. Schatz, LiPing Li, Quinn E. Carroll, Guofa Liu, Matthew J. Paul, Cecilia Flores

**Affiliations:** Department of Psychiatry, McGill University, Montréal, Quebec, Canada; Douglas Mental Health University Institute, Montréal, Quebec, Canada; Neuroscience Program, University at Buffalo, SUNY, New York, USA; Integrated Program in Neuroscience, McGill University, Montréal, QC, Canada; Department of Psychology, University at Buffalo, SUNY, New York, USA; Department of Biological Sciences, University of Toledo, Ohio, USA; Department of Neurology and Neurosurgery, McGill University, Montréal, Quebec, Canada; Ludmer Centre for Neuroinformatics & Mental Health, McGill University, Montréal, Quebec, Canada

**Keywords:** dopamine, prefrontal cortex, inhibitory control, axon guidance, puberty, sex differences

## Abstract

Dopamine axons are the only axons known to grow during adolescence. Here, using rodent models, we examined how two proteins, Netrin-1 and its receptor, UNC5C, guide dopamine axons towards the prefrontal cortex and shape behaviour. We demonstrate in mice (*Mus musculus*) that dopamine axons reach the cortex through a transient gradient of Netrin-1 expressing cells – disrupting this gradient reroutes axons away from their target. Using a seasonal model (Siberian hamsters; *Phodopus sungorus*) we find that mesocortical dopamine development can be regulated by a natural environmental cue (daylength) in a sexually dimorphic manner – delayed in males, but advanced in females. The timings of dopamine axon growth and UNC5C expression are always phase-locked. Adolescence is an ill-defined, transitional period; we pinpoint neurodevelopmental markers underlying this period.

## Introduction

Adolescence is a critical developmental period involving dramatic changes in behaviour and brain anatomy. The prefrontal cortex, the brain region responsible for our most complex cognitive functions, is still establishing connections during this time (Gogtay et al., 2004; Petanjek et al., 2011; Sowell et al., 2004). The trajectory of prefrontal cortex development in adolescence determines the vulnerability or resilience of individuals to adolescent-onset psychiatric diseases (Fuhrmann et al., 2015; Keshavan et al., 2014; Kessler et al., 2007, 2005; Lee et al., 2014). The age at which this adolescent development occurs therefore represents a critical window during which the brain is particularly susceptible to environmental influences. Traditionally, the onset of adolescence is thought to coincide with puberty (Hollenstein and Lougheed, 2013). In humans, the age of pubertal onset has been advancing throughout the 19^th^, 20^th^ and 21^st^ centuries, and environmental influences, such as nutrition, can pathologically alter the age of puberty (Wolf and Long, 2016). However, it remains entirely unknown whether the neural and cognitive maturational processes of adolescence can also be plastic. Here we examine how the timing of certain adolescent developmental processes are programmed, and whether this timing can be plastic in response to a natural environmental cue, in parallel with pubertal plasticity.

Dopamine innervation to the prefrontal cortex increases substantially across adolescence, and psychopathologies of adolescent origin prominently feature dopamine dysfunction. Evidence continues to emerge that protracted dopamine innervation is a key neural process underlying the cognitive and behavioural changes that characterize adolescence (Larsen and Luna, 2018). The mesocorticolimbic dopamine system – which includes the prefrontal cortex – is unique because not only are connections being formed and lost during adolescence, but there is also long-distance displacement of dopamine axons between brain regions. At the onset of adolescence, both mesolimbic and mesocortical dopamine axons innervate the nucleus accumbens in rodents, but the mesocortical axons leave the accumbens and grow towards the prefrontal cortex during adolescence and early adulthood (Hoops et al., 2018; Reynolds et al., 2018a). This is the only known case of axons growing from one brain region to another so late during development (Hoops and Flores, 2017).

The prolonged growth trajectory renders mesocortical dopamine axons particularly vulnerable to disruption. Environmental insults during adolescence (e.g. drug abuse) alter the extent and organization of dopamine innervation in the prefrontal cortex, leading to behavioural and cognitive changes in mice throughout adulthood (Drzewiecki and Juraska, 2020; Hoops and Flores, 2017; Reynolds and Flores, 2021). These changes often involve cognitive control, a prefrontal function that develops in parallel with dopamine innervation to the cortex in adolescence (Luna et al., 2015). Disruption of dopamine innervation frequently seems to result in “immature” cognitive control persisting through adulthood (Larsen and Luna, 2018).

Here, we examine the guidance of growing dopamine axons to the prefrontal cortex, and its timing. The guidance cue molecule Netrin-1, upon interacting with its receptor DCC, determines *which* dopamine axons establish connections in the nucleus accumbens and which ones leave this region to grow to the prefrontal cortex (Hoops and Flores, 2017; Reynolds and Flores, 2021). We hypothesized that the answers to *how* and *when* this extraordinary developmental feat is achieved may also lie in the Netrin-1 signalling system.

### Part 1: Netrin-1 “paves the way” for dopamine axons in adolescence

To identify the route by which dopamine axons grow from the nucleus accumbens to the medial prefrontal cortex, we visualized dopamine axons in the adult mouse forebrain. We observed that dopamine axons medial to the nucleus accumbens occupy a distinct area and are oriented dorsally towards the cortex (Figure 1A,B). Individual fibres can be seen crossing the boundary of the nucleus accumbens shell and joining these dorsally-oriented axons (Figure 1C). We hypothesized that these are the fibres that grow to the prefrontal cortex during adolescence. If this is correct, the number of dopamine axons oriented dorsally towards the medial prefrontal cortex should continue to increase until adulthood. To test this, we used a modified unbiased stereological approach (Kim et al., 2011) where axons are counted only if they crossed the upper and lower bounds of a counting probe. We also measured the average width of the area these axons occupy. We found, in both male and female mice, that the density of dopamine axons does not change between adolescence (21 days old) and adulthood (75 days old; Figure 1D). However, the width of the area that dopamine axons occupy does change, increasing between adolescence and adulthood (Figure 1E). These results indicate that the total number of dopamine axons passing through this area increases over adolescence and that dopamine axons grow to the medial prefrontal cortex via this route.

**Figure 1.**
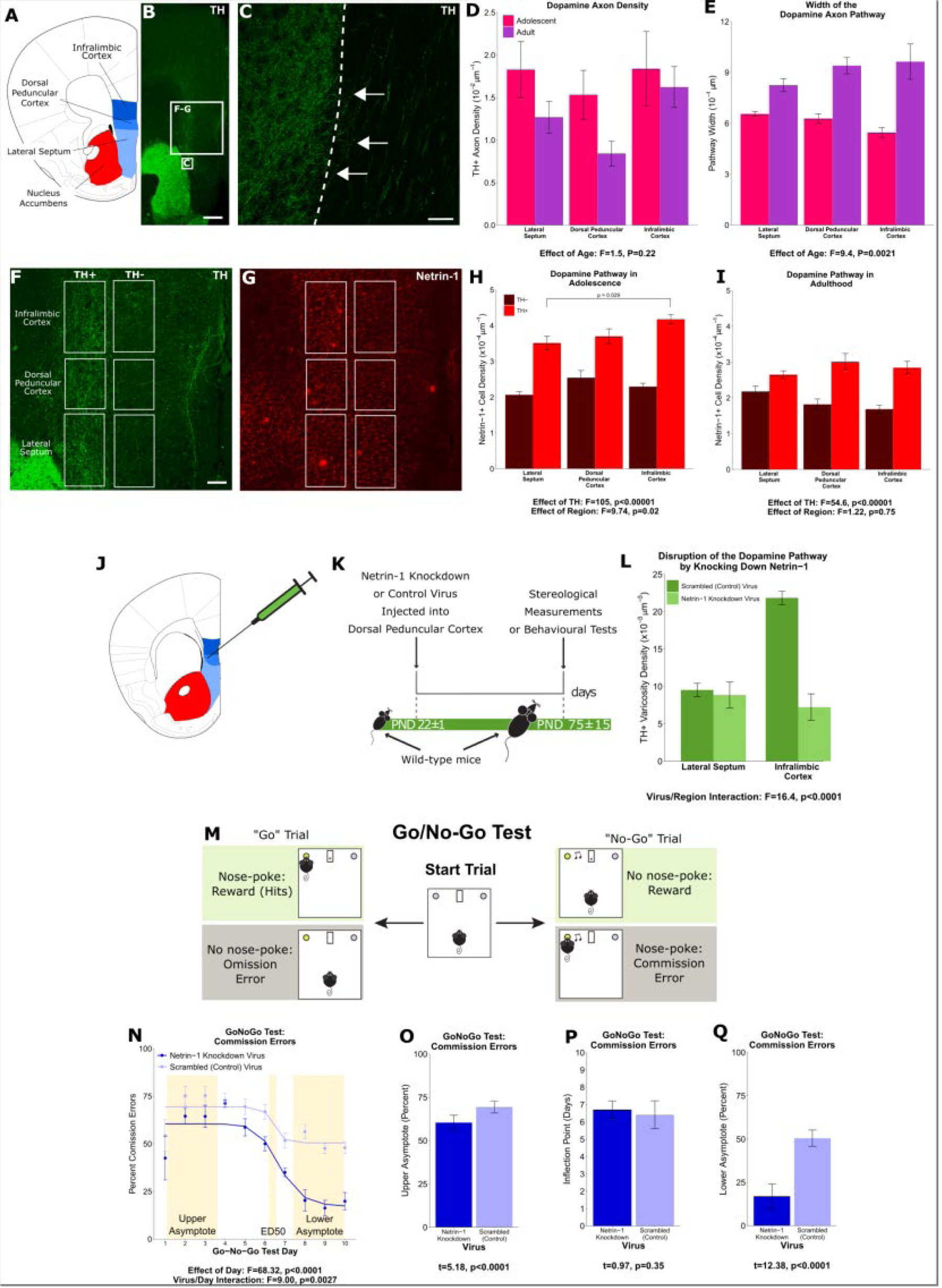
A “pathway” of Netrin-1 expressing cells “paves the way” for dopamine axons growing from the nucleus accumbens to the medial prefrontal cortex during adolescence. **A,** The brain regions containing the of dopamine fibres passing to the medial prefrontal cortex are highlighted in a line drawing of a coronal mouse brain section derived from Paxinos and Franklin (Paxinos and Franklin, 2013). **B,** An image of a coronal section through the forebrain of an adult mouse at low magnification (4x). Green fluorescence indicates immunostaining for tyrosine hydroxylase (TH), used here as a marker for dopamine. The smaller and larger white squares indicate the regions enlarged in panel C and panels F & G, respectively. Scale bar = 500 μm. **C,** The nucleus accumbens (left of the dotted line) is densely packed with TH+ axons (in green). Some of these TH+ axons can be observed extending from the nucleus accumbens medially towards TH+ fibres oriented dorsally towards the medial prefrontal cortex (white arrows). Scale bar = 10 μm. **D,** Modified stereological quantification revealed no significant difference in TH+ axon density between adolescence (21 days old) and adulthood (75 days old). Mixed-effects ANOVA, effect of age: F=1.53, p=0.22; region by age interaction: F=1.44, p=0.49. **E,** The average width of the area that dopamine axons occupy increased significantly from adolescence to adulthood, revealing that there is an increase in the total number of fibres passing to the medial prefrontal cortex during this period. Mixed-effects ANOVA, effect of age: F=9.45, p=0.0021; region by age interaction: F=5.74, p=0.057. **F,** In order to quantify the Netrin-1 positive cells along the TH+ fibre pathway, the pathway was contoured in each region, and a contour of equal area was placed medial to the dopamine pathway as a negative control. Scale bar = 200 μm. **G,** Using quantitative stereology, Netrin-1 positive cell density was determined along and adjacent to the pathway for each region. Red fluorescence indicates immunostaining for Netrin-1. **H,** In adolescent mice there are more Netrin-1 positive cells along the fibres expressing TH (“TH+”) than medial to them (“TH-“). This is what we refer to as the “Netrin-1 pathway”. Along the pathway, there is a significant increase in Netrin-1 positive cell density in regions closer to the medial prefrontal cortex, the innervation target. Mixed-effects ANOVA, effect of TH: F=105, p<0.0001. Effect of region: F=9.74, p=0.021. A post-hoc Tukey Test revealed a difference (p = 0.029) between the densities of the lateral septum and infralimbic cortex, but only within the dopamine pathway. **I,** In adult mice the Netrin-1 pathway is maintained, however there is no longer an increasing density of Netrin-1 positive cells towards the medial prefrontal cortex. Mixed-effects ANOVA, effect of TH: F=54.56, p<0.0001. Effect of region: F=1.22, p=0.75. **J,** The virus injection location within the mouse brain. A Netrin-1 knockdown virus, or a control virus, was injected into the dopamine pathway at the level of the dorsal peduncular cortex. **K,** Our experimental timeline: at the onset of adolescence a Netrin-1 knockdown virus, or a control virus, was injected in wild-type mice. In adulthood the mice were sacrificed and stereological measurements taken. **L,** TH+ varicosity density was quantified in the region below the injection site, the lateral septum, and in the region above the injection site, the infralimbic cortex. There was a significant decrease in TH+ varicosity density only in the infralimbic cortex. Mixed-effects ANOVA, virus by region interaction: F=16.41, p<0.0001. **M,** The experimental set-up of the final (test) stage of the Go/No-Go experiment. A mouse that has previously learned to nose-poke for a reward in response to a visual cue (illuminated nose-poke hole) must now inhibit this behaviour when the visual cue is paired with an auditory cue (acoustic tone). **N,** Mice injected with the Netrin-1 knockdown virus show improved action impulsivity compared to controls; they incur significantly fewer commission errors across the Go/No-Go task. Mixed-effects ANOVA, effect of day: F=68.32, p<0.0001. Day by virus interaction: F=9.00, p=0.0027. A sigmoidal curve is fit to each group of mice to determine how the two groups differ. Points indicate group means and error bars show standard error means. **O,** During the first days of Go/No-Go testing, both groups incur commission errors with high frequency, but the Netrin-1 knockdown group has fewer errors than the control group (T-test, t=5.18, p<0.0001). **P,** The ED50 – the inflection point in each sigmoidal curve – does not differ between groups, indicating that all mice improve their ability to inhibit their behavior at around the same time (T-test, t=0.97, p=0.35). **Q,** Mice microinfused with the Netrin-1 knockdown virus incur substantially fewer commission errors in the last days of the Go/No-Go task compared to mice injected with the control virus (T-test, t=12.38, p<0.0001). For all barplots, bars indicate group means and error bars show standard error means.

Next, we focussed on Netrin-1, a secreted protein that acts as a guidance cue to growing axons and is important for dopamine axon targeting in the nucleus accumbens in adolescence (Cuesta et al., 2020). Using unbiased stereology, we quantified the number of Netrin-1 expressing cell bodies along the dopamine axon route, and in an adjacent medial region as a control (Figure 1 F, G). We found that in adolescence there are more Netrin-1 positive neurons within the dopamine axon route than adjacent to it. Furthermore, along the axon route the density of Netrin-1 positive cells increases towards the medial prefrontal cortex, forming a dorsoventral gradient (Figure 1H). In adulthood, there remains a higher density of Netrin-1 positive cells along the dopamine route compared to the adjacent region, however the dorsoventral gradient is no longer present (Figure 1I).

To determine if Netrin-1 along the dopamine axon route is necessary for axon navigation, we silenced Netrin-1 expression in the dorsal peduncular cortex, the transition region between the septum and the medial prefrontal cortex, at the onset of adolescence (Figure 1J,K). In adulthood, we quantified the number of dopamine axon terminals in the regions below and above the injection site. Silencing Netrin-1 did not alter dopamine terminal density below the injection, in the lateral septum (Figure 1L). In the infralimbic cortex, which is the first prefrontal cortical region the axons reach after the injection site, terminal density was reduced in the Netrin-1 knock-down group compared to controls (Figure 1L). The knock-down appears to erase the Netrin-1 path to the prefrontal cortex, resulting in dopamine axons failing to reach their correct innervation target.

It remains unknown exactly what types of cells are expressing Netrin-1 along the dopamine axon route, and how this expression is regulated to produce the Netrin-1 gradients that guide the dopamine axons. It also remains unclear where the misrouted axons end up in adulthood. Future experiments aimed at addressing these questions will provide further valuable insight into the nature of the “Netrin-1 pathway”. Nonetheless, our results allow us to conclude that Netrin-1 expressing cells “pave the way” for dopamine axons growing to the medial prefrontal cortex.

We next examined how the Netrin-1 pathway may be important for behaviour. Dopamine input to the prefrontal cortex is a key factor in the transition from juvenile to adult behaviours that occurs in adolescence. We hypothesized that cognitive processes involving mesocortical dopamine function would be altered when these axons are misrouted in adolescence. To test our hypothesis, we used the Go/No-Go behavioural paradigm. This test quantifies inhibitory control, which matures in parallel with the innervation of dopamine axons to the prefrontal cortex in adolescence (Casey et al., 2008; Klune et al., 2021; Luna et al., 2015; Paus, 2005; Reynolds and Flores, 2021; Spear, 2000), and it is impaired in adolescent-onset disorders like depression and schizophrenia (Catts et al., 2013; Clementz et al., 2016; McTeague et al., 2016; Millan et al., 2012).

At the onset of adolescence, we injected the Netrin-1 silencing, or a scrambled control virus, bilaterally into the dorsal peduncular cortex; in adulthood we tested the mice in the Go/No-Go task. This paradigm first involves discrimination learning and reaction time training (Supplementary Figure 1a), followed by a Go/No-Go test consisting of “Go” trials where mice respond to a cue as previously trained and “No-Go” trials where mice must abstain from responding to the cue (Figure 1M). Correct responses to both trial types are reinforced with a food reward. We quantified the percent of “No-Go” trials where the mice incorrectly responded to the cue (“Commission Errors”) and the percent of “Go” trials where the mice correctly responded (“Rewards” or “Hits”; Supplementary Figure 1b). The ability of mice to respond correctly overall to both trial types is quantified as the Correct Response Rate (Supplementary Figure 1c) (Cuesta et al., 2019; Reynolds et al., 2018a, 2018b; Vassilev et al., 2021).

Mice injected with the Netrin-1 silencing virus differed from controls in their performance during “No-Go” trials. As the mice learn to withhold their responses over the course of the test, the number of commission errors they made in No-Go trials decreased in a sigmoidal fashion (Figure 1N). The upper and lower asymptotes of the sigmoidal curve quantify the number of commission errors committed during early and late test days, respectively, while the inflection point (ED50) indicates when mice start improving their ability to inhibit their behavior. At the start of the Go/No part of the task, the Netrin-1 silencing group make slightly fewer commission errors (Figure 1O) than control groups, although both groups begin to improve in the No-Go task at around the same time. However, the Netrin-1 silencing group achieved a substantially higher level of behavioural inhibition, quantified as a lower percentage of commission errors in the last test days (Figure 1Q), indicating an improved ability to withhold their behaviour on cue. These behavioural results demonstrate that the maturation of action impulsivity is sensitive to the organization of the ventro-dorsal Netrin-1 path that guides mesocortical dopamine axon growth. Deviations in this route associate with striking changes in the cognitive development that is characteristic of adolescence. In this case, the deviation leads to improved action impulsivity, suggesting that these dopamine axons may end up ectopically innervating a forebrain region other than the medial prefrontal cortex, enhancing cognitive control.

### Part 2: UNC5C expression coincides with the onset of adolescence

When axons leave the nucleus accumbens during adolescence, they follow a Netrin-1 “path” through intermediate brain regions to reach their intended innervation target. However, only a small subset of the dopamine axons that have reached the nucleus accumbens by early adolescence leave; the vast majority stay and form connections in the accumbens throughout life (Reynolds et al., 2018a). The “decision making” process of whether to “stay” (in the accumbens) or “go” (to the cortex via the Netrin-1 path) happens during a narrow developmental window at the onset of adolescence (Reynolds et al., 2018b). It remains unknown how the timing of this process is determined.

**Figure 2.**
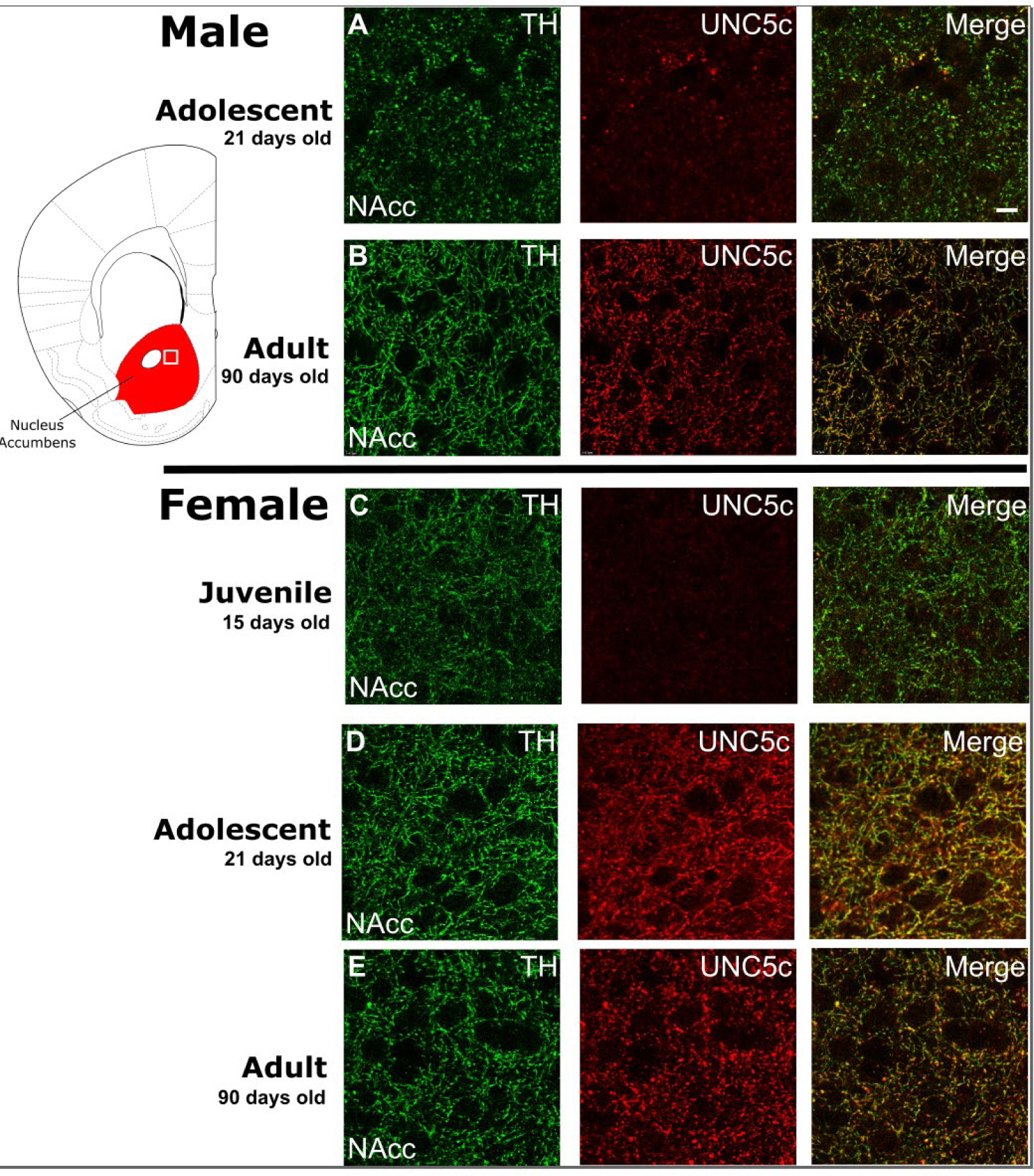
The age of onset of UNC5C expression by dopamine axons in the nucleus accumbens of mice is sexually dimorphic. Images are representative of observed immunofluorescence patterns in the nucleus accumbens (approx. location highlighted as a white square in the coronal mouse brain section Plate 19, modified from Paxinos & Franklin, 2013). No qualitative differences were noted between the shell and core of the nucleus accumbens. For each row, six individuals were sampled. In males (**A-B**), UNC5C expression on dopamine fibres (here identified by immunofluorescent staining for tyrosine hydroxylase, TH) in the nucleus accumbens appears during adolescence. **A,** At the onset of adolescence (21 days old) dopamine fibres do not express UNC5C. Scale bar = 10 μm. **B,** By adulthood (90 days old), dopamine fibres express UNC5C. In females (**C-E**), UNC5C expression on dopamine fibres in the nucleus accumbens appears prior to adolescence. **C,** In juvenile (15 day old) mice, there is no UNC5C expression on dopamine fibres. **D,** By adolescence, dopamine fibres express UNC5C. **E,** In adulthood, dopamine fibres continue to express UNC5C.

In adolescence, dopamine neurons begin to express the repulsive Netrin-1 receptor UNC5C, particularly when mesolimbic and mesocortical dopamine projections segregate in the nucleus accumbens (Manitt et al., 2010; Reynolds et al., 2018a). In contrast, dopamine axons in the prefrontal cortex do not express UNC5c, except in very rare cases (Supplementary Figure 2a). In adult male mice with *Unc5c* haploinsufficiency, there appears to be ectopic growth of mesolimbic dopamine axons to the prefrontal cortex (Auger et al., 2013). This miswiring is associated with alterations in prefrontal cortex-dependent behaviours (Auger et al., 2013).

Using immunohistochemistry, we assessed the expression of UNC5C on nucleus accumbens dopamine axons across development. In male mice, we found little expression of UNC5C on dopamine axons at the onset of adolescence (Figure 2A), while we did find UNC5C expression on dopamine axons in adults (Figure 2B). Remarkably, when we assessed this in females, we found dopamine axons already expressing UNC5C in the nucleus accumbens at the onset of adolescence (Figure 2D), similar to adult females (figure 2E), indicating that the onset of UNC5C expression on dopamine axons in the nucleus accumbens is sexually dimorphic, with an earlier emergence in females. We examined the nucleus accumbens in pre-adolescent female mice and indeed found little UNC5C expression on dopamine axons (Figure 2C). The onset of UNC5C expression in mesocorticolimbic dopamine axons is therefore peri-adolescent but occurs earlier in females than in males, consistent with the earlier emergence of adolescence in female rodents and the earlier onset of adolescence and puberty in humans (Wolf and Long, 2016). Differences in the precise timing of dopamine innervation to the PFC in adolescence have been suggested by findings reported in male and female rats (Willing et al., 2017)”.

### Part 3: Environmental control of the timing of adolescence

We hypothesize that at the emergence of adolescence, UNC5C expression by dopamine axons in the nucleus accumbens signals the initiation of the growth of dopamine axons to the prefrontal cortex. We therefore examined whether the developmental timings of UNC5C expression and dopamine innervation of the prefrontal cortex are similarly affected by an environmental cue known to delay pubertal development in seasonal species.

Siberian hamsters (*Phodopus sungorus*) regulate many aspects of their behavior and physiology to meet the changing environmental demands of seasonality (Paul et al., 2008; Stevenson et al., 2017). In winter, they increase the thickness of their fur, exchange their brown summer coats for white winter ones, and undergo a daily torpor to conserve energy (Scherbarth and Steinlechner, 2010). In addition, adults suppress reproduction and juveniles delay puberty (Pevet, 1988; Yellon and Goldman, 1984), including developmental changes in gonadotropin releasing hormone neurons in the hypothalamus (Buchanan and Yellon, 1991; Heywood and Yellon, 1997). Reproductive organ development is delayed as part of pubertal postponement (Darrow et al., 1980; Ebling, 1994; Timonin et al., 2006). This seasonal plasticity is regulated by long or short periods of daylight (Heldmaier and Steinlechner, 1981; Hoffmann, 1978) and raises the possibility that aspects of adolescent development are sensitive to these environmental cues. To our knowledge, adaptive variation in the timing of adolescent neural development has never been recorded in any animal.

Here, we tested whether day length regulates *when* dopamine axons grow to the cortex, and whether the timing of UNC5C expression in the nucleus accumbens and adolescent changes in behavior are similarly affected.

### 3.i The seasonality of adolescence

Male hamsters were examined at three ages: 15 days old (±1), 80 days old (±10), and 215 days old (±20). We compared the density of the dopamine innervation to the medial prefrontal cortex in male hamsters housed under lighting conditions that replicate summer daylengths (long days, short nights) or winter daylengths (short days, long nights) (Figure 3A,B). We will refer to these two groups as “summer hamsters” and “winter hamsters” to emphasize the natural stimulus we are replicating in the laboratory environment. We confirmed that puberty is delayed in male winter hamsters compared to summer hamsters in the present experiment by measuring their gonadal weights across ages (Supplementary Figure 3a).

In male summer hamsters, dopamine input density to the prefrontal cortex increases during adolescence, after 15 days old and before 80 days old (Figure 3C), consistent with dopamine axon growth in mice (Manitt et al., 2013, 2011; Reynolds et al., 2018a). Prefrontal cortex dopamine innervation in summer hamsters continues to increase after 80 days old (Figure 3C).

In male winter hamsters, dopamine innervation to the prefrontal cortex is delayed until after 80 days, which coincides with their delayed pubertal onset (Figure 3D, Supplementary Figure 3a). This demonstrates that an environmental cue can determine the timing of adolescent brain development.

**Figure 3.**
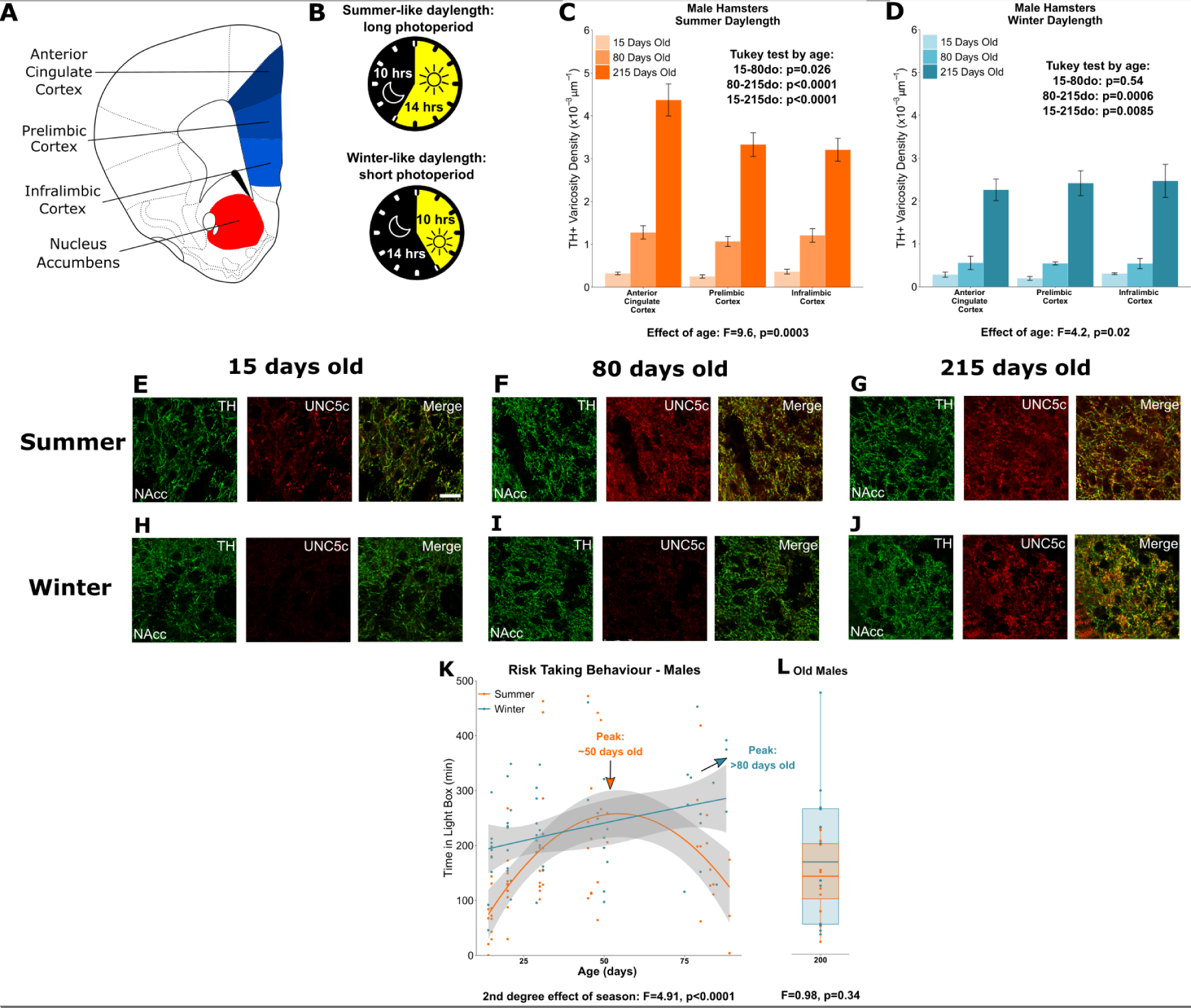
Plasticity of adolescent development in male Siberian hamsters according to seasonal phenotype. All results illustrated in this figure refer to results in male hamsters. **A,** Dopamine innervation was quantified in three subregions of the medial prefrontal cortex, highlighted in blue. UNC5C expression was examined in the nucleus accumbens, highlighted in red. Line drawing of a coronal section of the mouse brain was derived from Plate 19 of Paxinos and Franklin (Paxinos and Franklin, 2013). **B,** Hamsters were housed under either summer-mimicking long days and short nights (“summer hamsters”) or winter-mimicking short days and long nights (“winter hamsters”). **C,** In male hamsters housed under a summer-mimicking daylength there is an increase in dopamine varicosity density in the medial prefrontal cortex between 15 and 80 days old. Mixed-effects ANOVA, effect of age: F=9.6, p=0.000255. Tukey Test, 15-80 days old (do): p=0.026; 80-215do: p<0.0001; 15-215do: p<0.0001. **D,** In male hamsters housed under a winter-mimicking daylength there is no increase in dopamine varicosity density until hamsters have reached 215 days of age. Mixed-effects ANOVA, effect of age: F=4.17, p=0.0205. Tukey Test, 15-80do: p=0.54; 80-215do: p=0.0006; 15-215do: p=0.0085. **E,** At 15 days old, dopamine axons (here identified by immunofluorescent staining for tyrosine hydroxylase, TH) in the nucleus accumbens of male summer-daylength hamsters largely do not express UNC5C. Scale bar = 20um (bottom right). **F-G,** At 80 (**F**) and 215 (**G**) days old, dopamine axons in the nucleus accumbens express UNC5C. **H-I,** At 15 (**H**) and 80 (**I**) days old, dopamine axons in the nucleus accumbens of male winter hamsters largely do not express UNC5C. **J,** By 215 days old there is UNC5C expression in dopamine axons in the nucleus accumbens of male winter hamsters. **E-J**, Representative images of the nucleus accumbens shell, 6 individuals were examined per group. **K,** Male hamsters house under a summer-mimicking daylength show an adolescent peak in risk taking in the light/dark box apparatus. Those raised under a winter-mimicking photoperiod show a steady increase in risk taking over the same age range. Arrows indicate the ages at which risk taking peaks in summer (orange) and winter (blue) hamsters. Polynomial regression, effect of season: F=3.551, p=0.00056. **L,** In male hamsters, at 215 days of age, there is no difference in risk taking between hamsters raised under summer and winter photoperiods. T-test, effect of season: t=0.975, p=0.341. For all barplots, bars indicate group means and error bars show standard error means.

We then examined UNC5C expression by dopamine axons in the nucleus accumbens in male summer and winter hamsters across age classes. UNC5C expression was apparent only after the onset of adolescence in summer hamsters (Figure 3E,F,G), as observed in male mice. However, UNC5C expression was delayed in male winter hamsters – this group did not show UNC5C expression in dopamine axons in the nucleus accumbens until after 80 days old (Figure 3H,I,J). This aligns with the delayed timing of mesocortical dopamine axon growth and pubertal onset in male winter hamsters and demonstrates that the emergence of UNC5C is a marker of adolescent onset in male mice.

A behavioural characteristic of adolescence is increased willingness to enter a novel environment, a behaviour that assumes an increased amount of risk (Arrant et al., 2013; Lynn and Brown, 2009). To measure this, we used the light/dark test (Bourin and Hascoët, 2003). Time spent in the light compartment is dopamine-dependent (Bahi and Dreyer, 2019; Gao and Cutler, 1993) and peaks in adolescence (Arrant et al., 2013). We will refer to this behaviour as “risk taking”. We assessed the developmental profile of risk taking in the light/dark box test in summer and winter hamsters across adolescence. In male summer hamsters, the risk taking increases across adolescence, peaks around 50 days, then subsequently declines (Figure 3K). However, the adolescent increase in risk taking is protracted in winter hamsters: across the age range examined we observe a gradual, consistent increase in risk taking rather than a peak and decline.

We next assessed a cohort of 215-day old hamsters, for which both summer and winter male hamsters have undergone puberty and exhibit high levels of dopamine innervation of the prefrontal cortex (Figure 3C,D,G,J, Supplementary Figure 3a). In these hamsters, we find no difference in risk taking between the male summer and winter groups (Figure 3L), demonstrating that, after 80 days, risk taking begins to decline in male winter hamsters and that by 215 days it has declined to the same level as in summer hamsters. Male hamsters raised under summer-mimicking long days and winter-mimicking short days both ultimately make the transition to the adult behavioral phenotype.

### 3.ii An extraordinary case of decoupling puberty and adolescence

In parallel with males, we conducted equivalent experiments in female hamsters (Figure 4A,B). Under a summer-mimicking daylength, dopamine innervation to the medial prefrontal cortex increases between 15 and 80 days old, similar to male summer hamsters (Figure 4C). There is no further increase in innervation density after 80 days old, consistent with earlier adolescent development in females observed in other rodent species (Juraska and Willing, 2017; Kopec et al., 2018; Reynolds and Flores, 2021; Spear, 2000; Westbrook et al., 2018). We confirmed that puberty is delayed in female winter hamsters compared to summer hamsters by measuring their uterine weights (Supplementary Figure 4a) and vaginal opening (Supplementary Figure 4b) across ages.

**Figure 4.**
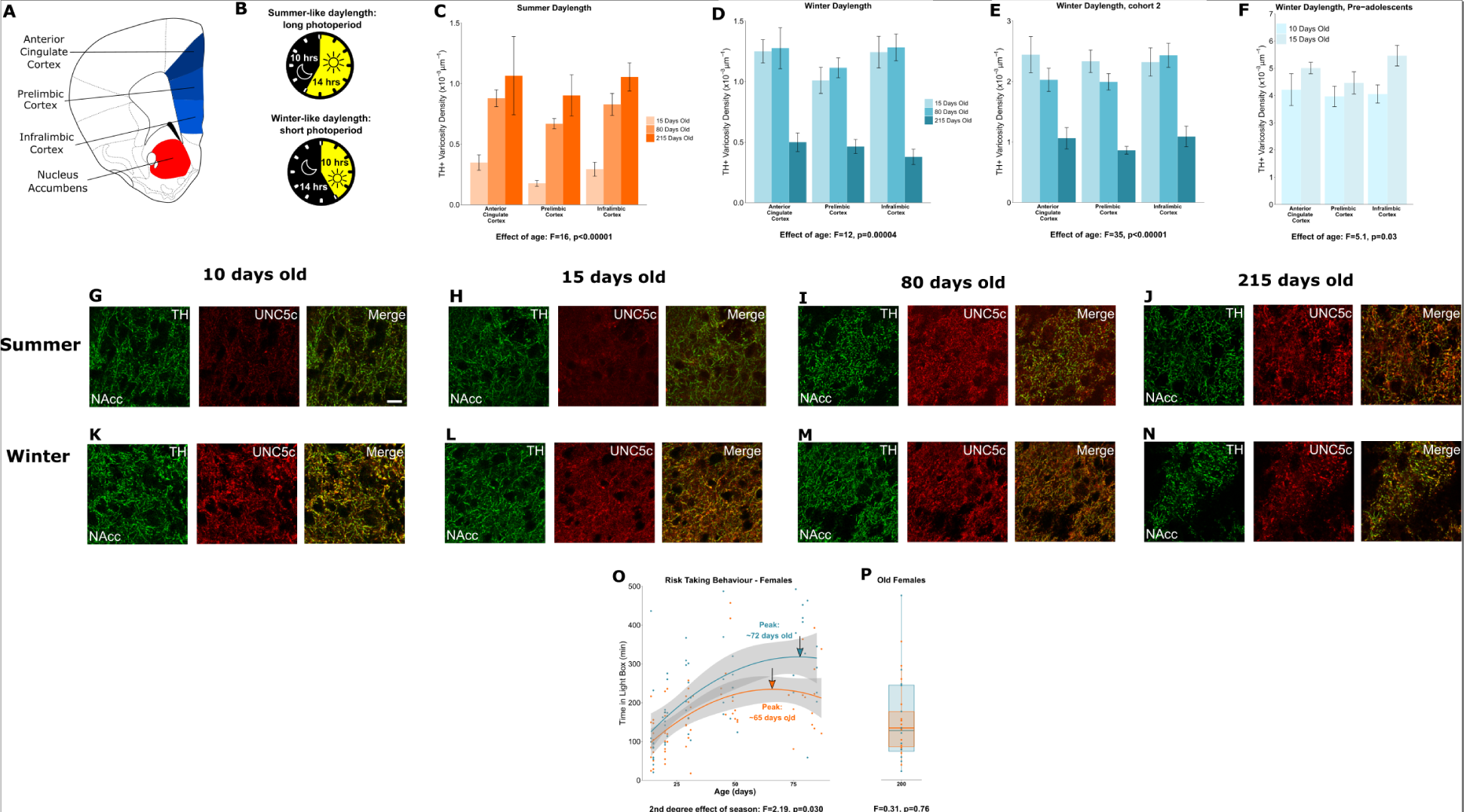
Plasticity of adolescent development in female Siberian hamsters according to seasonal phenotype. All results illustrated in this figure refer to results in female hamsters. **A,** Dopamine innervation was quantified in three subregions of the medial prefrontal cortex, highlighted here in blue. UNC5C expression was examined in the nucleus accumbens, highlighted in red. Line drawing of a coronal section of the mouse brain was derived from Paxinos and Franklin (Paxinos and Franklin, 2013). **B,** Hamsters were housed under either a summer-mimicking or winter-mimicking daylength. **C,** In female hamsters housed under a summer daylength dopamine varicosity density in the medial prefrontal cortex increases between 15 and 80 days of age. Mixed-effects ANOVA, effect of age: F=16.72, p<0.0001 **D,** In female hamsters housed under a winter daylength there is no increase in dopamine varicosity density post-adolescence. Instead, there is a steep decline in density between 80 and 215 days of age. Mixed-effects ANOVA, effect of age: F=12.33, p=0.000043. **E,** As our results in panel D were unexpected, we replicated them with a second cohort of hamsters and found qualitatively identical results. Mixed-effects ANOVA, effect of age: F=34.871, p<0.0001. **F,** To try and determine when dopamine varicosities innervate the medial prefrontal cortex, we examined a cohort of 10- and 15-day-old hamsters. We found that varicosity density increases in the medial prefrontal cortex during this time, indicating that dopamine innervation to the medial prefrontal cortex is accelerated in female winter hamsters. Mixed-effects ANOVA, effect of age: F=5.05, p=0.03. **G-H,** In 10- and 15-day-old female summer hamsters there is little UNC5C expression in nucleus accumbens dopamine axons (here identified by immunofluorescent staining for tyrosine hydroxylase, TH). Sample size: 4 (panel G) or 6 (panel H). **I-J,** By 80 days old (panel I), and continuing at 215 days old (panel J), dopamine axons in the nucleus accumbens express UNC5C in female summer hamsters. Sample sizes: 6. Scale bar = 20um (panel G bottom right). **K-N,** At all ages which winter female hamsters were examined, dopamine axons in the nucleus accumbens express UNC5C in winter female hamsters. Sample sizes: 4 (panel K) or 6 (panels L-N). **O,** In female hamsters, those raised under summer and winter daylengths both show an increase in risk taking over time. The winter hamsters peak later compared to the summer daylength hamsters. Arrows indicate the ages at which risk taking peaks in summer (orange) and winter (blue) hamsters. Polynomial regression, effect of season: F=3.305, p=0.00126. **P,** In female hamsters, at 215 days of age, there is no difference in risk taking between hamsters raised under summer and winter photoperiods. T-test, effect of season: t=0.309, p=0.76. For all barplots, bars indicate group means and error bars show standard error means.

When housed under a winter-mimicking daylength, dopamine input density in the prefrontal cortex of female hamsters is *not* delayed as in males, but rather reaches adult levels prior to 15 days old (Figure 4D). We replicated this unexpected finding in a separate, independent cohort of female winter hamsters (Figure 4E). This surprising result shows an intervention that accelerates adolescent cortical development.

We then measured dopamine axon density in female winter hamsters at two earlier ages: 10 and 15 days old. Dopamine innervation increases during this period (figure 4F), well before normal adolescence and long before pubertal development. This is an extraordinary phenomenon: a key marker of adolescent neurodevelopment is accelerated and dissociated from puberty in female hamsters raised under winter-mimicking short days (Supplementary Figure 4a,4b).

The early increase in prefrontal cortex dopamine terminals in winter females is followed by a dramatic reduction between 80 and 215 days old (Figure 4D,E). This overlaps with the delayed timing of puberty in these females (Butler et al., 2007; Supplementary Figure 4a,4b). Synaptic pruning in the cortex is a well-known component of adolescent neural development across species (Huttenlocher, 1984; Koss et al., 2013; Petanjek et al., 2011). Under normal conditions, the effect of pruning on dopamine synapses is likely masked by the growth of new dopamine axons to the prefrontal cortex (Manitt et al., 2013, 2011; Reynolds et al., 2018a). In the case of female winter hamsters, we hypothesize that the growth of dopamine axons to the prefrontal cortex occurs early while synaptic pruning, including of dopamine synapses, appears to occur later. This leads to a remarkable dissociation between two cortical developmental processes that are normally simultaneous, the behavioural implications of which are unclear.

If the developmental onset of UNC5C expression determines the timing of dopamine innervation of the prefrontal cortex, then onset of UNC5C expression should also be advanced in female winter hamsters. Hence, we examined UNC5C expression at the same ages as we examined dopamine axon growth in female hamsters. At 10 and 15 days old, UNC5C expression is present *only* in the winter hamsters (Figure 4G,H,K,L), but at 80 and 215 days old, UNC5C expression is apparent in both summer and winter hamsters (Figure 4I,J,M,N).

We used the light/dark box test to examine potential risk taking implications of the extraordinary developmental trajectory we observed in the prefrontal cortex of female hamsters. In female summer and winter hamsters, the adolescent increase and peak in risk taking occurs between the ages of 15 and 80 days, as it does in summer daylength males (Figure 4O). However, contrary to what we would expect, the peak in winter females is delayed compared to summer females. When we assessed an independent cohort of 215-day-old female hamsters, we found no difference in risk taking between groups (Figure 4R), indicating that, like males, female summer and winter hamsters both eventually reach the same adult level of risk taking.

In both sexes, hamsters housed under a summer-mimicking daylength showed an adolescent peak in risk taking at an age that we would predict based on results from other rodents (Arrant et al., 2013; Pietropaolo et al., 2004; Tanaka, 2015). When raised under a winter-mimicking daylength, hamsters of either sex show a protracted peak in risk taking. In males, it is delayed beyond 80 days old, but the delay is substantially less in females. This is a counterintuitive finding considering that dopamine development in winter females appears to be accelerated. Our interpretation of this finding is that the timing of the risk taking peak in females may reflect a balance between different adolescent developmental processes. The fact that dopamine axon growth is accelerated does not imply that all adolescent maturational processes are accelerated. Some may be delayed, for example those that induce axon pruning in the cortex.

The timing of the risk taking peak in winter female hamsters may therefore reflect the amalgamation of developmental processes that are advanced with those that are delayed – producing a behavioural effect that is timed somewhere in the middle. Disentangling the effects of different developmental processes on behaviour will require further experiments in hamsters, including the direct manipulation of dopamine activity in the nucleus accumbens and prefrontal cortex.

## Conclusion

Here we describe how the gradual growth of mesocortical dopamine axons marks adolescent development, and how this process uses guidance cues and is sensitive to sex and environment. Netrin-1 signalling provides the “stay-or-go” “decision making” conducted by dopamine axons that innervate the nucleus accumbens at the onset of adolescence (Cuesta et al., 2020). UNC5C expression by these dopamine axons marks the timing at which this decision is made. In mice, UNC5C expression coincides with sex differences in both adolescent and pubertal development. Females, which develop earlier, show earlier UNC5C expression in dopamine axons compared to males.

In hamsters, behavioural and developmental shifts in response to environmental cues occur in parallel with alterations in the timing of dopamine axon growth. As we show here, male hamsters raised under a winter-mimicking daylength delay not only puberty, but also adolescent dopamine and behavioural maturation. In contrast, female hamsters under identical conditions delay puberty but accelerate dopamine axon growth, a key marker of adolescent brain development. Behavioural shifts during adolescence appear to be delayed in these females, but less substantially than in male hamsters. Notably, under all conditions, the developmental timing of UNC5C expression corresponds to the timing of dopamine innervation of the prefrontal cortex.

In both mice and hamsters, the emergence of UNC5C expression coincides with the onset of dopamine axon growth to the prefrontal cortex, a key characteristic of the adolescent transition period. While previously we have shown that the Netrin-1 signalling in the nucleus accumbens is responsible for coordinating *whether* dopamine axons grow in adolescence (Reynolds and Flores, 2021), here we propose that Netrin-1 signalling is also key to determining *how* and *when* this marker of adolescence occurs.

## Supporting information

Supplementary Statistics

## Acknowledgments

We are grateful for the technical assistance of the Pathology Core, Centre for Phenogenomics, Toronto, Canada, and in particular Milan Ganguly and Gregory Ossetchkine, for their excellent histological assistance. Furthermore, we are grateful to Helen Cooper and Philip Vassilev for their critical and insightful readings of our manuscript.

## Author contributions (Brand et al., 2015)

Conceptualization: DH, MJP, CF

Data Curation: DH

Formal Analysis: DH

Investigation: DH, RFK, SS, EE, AH, TO, AD, CP, JZ, KCS, LPL, QEC

Investigation (post-review experiments): DM, RGA

Methodology: DH, MJP, CF

Resources: GL, MJP, CF Supervision: DH, MJP, CF

Validation: DH Visualization: DH

Visualization (post-review experiments): DH, DM, RGA

Writing—original draft: DH

Writing—review & editing: DH, MJP, CF

## Funding

National Institute on Drug Abuse grant R01DA037911 (CF) Canadian Institutes of Health Research

FRN: 156272 (CF) Canadian Institutes of Health Research

FRN: 170130 (CF)

National Science and Engineering Research Council of Canada RGPIN-2020-04703 (CF)

National Science Foundation IOS-1754878 (MJP)

National Science and Engineering Research Council of Canada PGSD3-415253-2012 (DH) Quebec

Nature and Technology Research Fund 208332 (DH)

National Science and Engineering Research Council of Canada PDF5171462018 (DH)

McGill-Douglas Max Planck Institute of Psychiatry International Collaborative Initiative in Adversity and Mental Health, an international partnership funded by the Healthy Brains for Healthy Lives initiative (RGA)

## Data and code availability

All data and code use in these analyses are available through the Open Science Framework (DIO 10.17605/OSF.IO/DU3H4).

