## Supplementary material for "The scheduling of adolescence with Netrin-1 and UNC5C": Methods Supplement

**Supplementary Documents: Methods**

**Animals**

All mouse (*Mus musculus*) experiments and procedures were performed in accordance with the guidelines of the Canadian Council of Animal Care and the McGill University/Douglas Hospital Research Centre Animal Care Committee. All mice were received from Charles River Canada and housed with same-sex littermates on a 12h light/dark cycle with ad libitum access to food and water. We used male mice for these experiments.

All Siberian hamster (*Phodopus sungorus*) experiments and procedures were approved by the University at Buffalo, SUNY Institutional Animal Care and Use Committee. Hamsters were obtained from our colony (MJP), which was originally derived from animals generously provided by Dr. Brian Prendergast, University of Chicago, in 2015. Hamsters were housed with same-sex littermates in well-ventilated, light-proof environmental housing units that provided either a summer-mimicking long-day photoperiod (14:10 hour light:dark cycle) or a winter-mimicking short-day photoperiod (10:14 hour light:dark cycle); dim red light was present during the dark phase. Food and water were available ad libitum. Both male and female mice were used for these experiments.

There is no universally accepted way to define the precise onset of adolescence. Therefore, there is no clear-cut boundary to define adolescent onset in rodents (Spear, 2000). Puberty can be more sharply defined, and puberty and adolescence overlap in time, but the terms are not interchangeable. Puberty is the onset of sexual maturation, while adolescence is a more diffuse period marked by the gradual transition from a juvenile state to independence. We, and others, suggest that adolescence in rodents spans from weaning (postnatal day 21) until adulthood, which we take to start on postnatal day 60 (Reynolds and Flores, 2021). We refer to "early adolescence" as the first two weeks postweaning (postnatal days 21-34). These ranges encompass discrete DA developmental periods (Kalsbeek et al., 1988; Manitt et al., 2011; Reynolds et al., 2018a), vulnerability to drug effects on DA circuitry (Hammerslag and Gulley, 2014; Reynolds et al., 2018a), and distinct behavioral characteristics (Adriani and Laviola, 2004; Makinodan et al., 2012; Schneider, 2013; Wheeler et al., 2013).

**Tissue Processing**

Rodents were euthanized with an intraperitoneal injection of 50 mg/kg ketamine, 5 mg/kg xylazine, and 1 mg/kg acepromazine. They were then perfused intracardially with 10 IU/mL heparinized saline (mice) or physiological saline (hamsters) followed by fixative solution (4% paraformaldehyde). Both perfused solutions were pH-adjusted to between 7.2-7.4 with dilute hydrochloric acid and sodium hydroxide. After perfusion, brains were dissected from the skull, placed in fixative solution overnight at 4°C and then stored in phosphate-buffered saline at 4°C. Brains were cut coronally into 30 um (hamster) or 35 um (mouse) thick sections on a vibratome.

**Immunohistochemistry**

Every second section (mouse) or third section (hamster) was processed for immunofluorescence as we have described previously (Salameh et al., 2017).

For experiments in mouse tissue requiring only TH staining, we used a rabbit anti-tyrosine hydroxylase (TH; 1:1000 dilution, product #AB152; Millipore) antibody as the primary antibody and an Alexa Fluor (AF) 594-conjugated donkey anti-rabbit antibody (1:500 dilution, product

#711585152, Jackson Laboratories) as the secondary antibody. We and others have shown that the TH antibody used in these studies labels dopamine axons but rarely labels norepinephrine axons within the regions of interest (Manitt et al., 2013, 2011; Miner et al., 2003; Reynolds et al., 2022).

To examine hamster sections for TH only, a 3,3'-Diaminobenzidine (DAB) staining protocol was used. First, we performed antigen retrieval using heated (70°C) Citrate Buffer (0.05M) followed by Glycine (0.1M). We used a mouse anti-TH antibody (1:22000 dilution, product #T1299, Sigma), followed by secondary staining using a DAB staining kit (product #SK4100, Vector Laboratories).

To detect both TH and Netrin-1, we used a mouse anti-TH antibody (1:1000 dilution, product #MAB318; Millipore) and a rabbit Netrin-1 antibody (1:500 dilution, product #ab126729, abcam) as primary antibodies. We use citrate buffer and sodium dodecyl substrate antigen recovery methods to strengthen the Netrin-1 signal as previously described (Salameh et al., 2017). We used AF488-conjugated donkey anti-mouse antibody (1:500 dilution, product #715545150, Jackson Laboratories) and the AF594-conjugated donkey anti-rabbit antibody (as above) as secondary antibodies.

To detect both TH and UNC5C, we first used the rabbit anti-TH antibody (1:1000 dilution) and a mouse anti-UNC5C antibody (1:100 dilution, provided by Dr. Guofa Liu) as primary antibodies and a AF488-conjugated donkey anti-rabbit (1:500 dilution, product #711545152, Jackson Laboratories) and AF594-conjugated donkey anti-mouse (1:500 dilution, product #711585152, Jackson Laboratories) secondary antibodies. For these experiments we used the same antigen retrieval methods as with our immunohistochemistry staining for Netrin-1 described above and in (Salameh et al., 2017).

To replicate our results with a commercially available antibody, we used a goat anti-UNC5C antibody (1:200 dilution, product #NBP1-37002, NOVUS Biologicals) along with the rabbit anti-TH antibody (1:500 dilution) as primary antibodies. We used the AF488-conjugated donkey anti-rabbit (as above) and AF594-conjugated donkey anti-goat (1:500 dilution, product #705585003, Jackson Laboratories) antibodies as secondary antibodies. For this experiment we used two variations on the standard immunohistochemistry protocol described in Salameh et al., 2017: tris-buffered saline was used in place of phosphate-buffered saline, and commercial protein block and antibody diluent (both from Agilent) were used in place of a bovine serum albumin blocking solution.

In all immunochemistry experiments we stain for TH as a marker for dopamine in order to identify dopamine axons. Therefore, we pay great attention to the morphology and localization of the fibres to avoid including in our study any fibres stained with TH antibodies that are not dopamine fibres. The fibres that we examine and that are labelled by the TH antibody show features indistinguishable from the classic features of cortical dopamine axons in rodents (Berger et al., 1983, 1974; Eden et al., 1987; Manitt et al., 2011), namely they are thin fibres with irregularly-spaced varicosities, are densely packed in the nucleus accumbens, sparsely present only in the deep layers of the prefrontal cortex, and are not regularly oriented in relation to the pial surface. This is in contrast to rodent norepinephrine fibres, which are smooth or beaded in appearance, relatively thick with regularly spaced varicosities, increase in density towards the shallow cortical layers, and are in large part oriented either parallel or perpendicular to the pial surface (Berger et al., 1983, 1974; Levitt and Moore, 1979; Miner et al., 2003). Furthermore, previous studies in rodents have noted that only norepinephrine cell bodies are detectable using immunofluorescence for TH, not norepinephrine processes (Miner et al., 2003; Pickel et al., 1975; Verney et al., 1982), and we did not observe any norepinephrine-like fibres. Finally, a DAT-Cre approach has been used to demonstrate that all axons that immunostain for TH in the forebrain are dopamine axons (Caldwell et al., 2023). We are not aware of any other processes in the forebrain that are known to be immunopositive for TH under any environmental conditions.

After immunofluorescence staining, sections were mounted on gel-coated slides and cover-slipped with a fluorescence-preserving mounting medium (“Vectashield” branded media, Vector Laboratories). Sections were either stained with DAPI prior to mounting or mounted with a DAPI-containing medium.

**Stereological Analyses**

For all experiments, contours were delineated on sections corresponding to plates 13-22 of the mouse brain atlas (Paxinos and Franklin, 2013) or plates 10-14 of the hamster brain atlas (Morin and Wood, 2001). The brain regions along the dopamine axon route from the nucleus accumbens to the prefrontal cortex consist of the lateral septum, dorsal peduncular cortex, and infralimbic cortex; the latter being the first medial prefrontal cortex subregion encountered along this route. The subregions of the medial prefrontal cortex in which dopamine varicosities were quantified in this study are the infralimbic cortex, prelimbic cortex, and anterior cingulate cortex. All regions were examined only anterior to the genu of the corpus callosum. Counting was conducted bilaterally in mice and in the left hemisphere in hamsters.

Dopamine axon density along the route from the nucleus accumbens to the medial prefrontal cortex was determined using a modified stereological approach based on that described in (Kim et al., 2011). The dense bundle of dopamine fibres that occurs along the lateral boundary of each region was traced at x 5 magnification with a Leica DM400B microscope and StereoInvestigator (Microbrightfield) software. Using the counting probe function of StereoInvestigator, a grid of 175 $um^2$ was superimposed on each contour, starting at a random starting point within the contour. Unbiased counting frames (length = 25 um, width = 10 um) were placed in the top left corner of each grid square. Axons were counted if they crossed both the upper and lower boundaries of the counting frame. Counting was conducted at x40 magnification using a counting depth of 10 um and a guard zone of 5 um. Counts were performed blind by a single individual (TO). Axon density was determined by dividing the total axon count by the width of the contour.

Netrin-1-positive cell bodies were used as the counting unit to examine Netrin-1 density along the dopamine axon route from the nucleus accumbens to the medial prefrontal cortex. The dense bundle of dopamine fibres that occurs along the lateral boundary of each region was traced at x5 magnification with a Leica DM400B microscope and StereoInvestigator software. We also delineated a region of equal area directly medial to the fiber bundle, and we considered this the TH-negative subregion (Figure 1F of the main text). To determine the number of Netrin-1-positive cell bodies, we used the optical fractionator probe function of Stereoinvestigator with a grid of 175 $um^2$, an unbiased counting frame of 100 $um^2$, a counting depth of 10 um, and a guard zone of 2 um. Counting was conducted at x40 magnification using the standard counting protocol for quantifying discrete objects (“particle stereology”;  (Howard and Reed, 2004). Counts were performed blind by a single individual (SS). To determine the volume of each subregion we used the Cavalieri method in Stereoinvestigator (Howard and Reed, 2004). The coefficient of error was below 0.1 for all measures. Cell density was determined by dividing the total count of cells by the volume of the subregion.

TH-positive varicosities were used as the counting unit to obtain a measure of dopamine presynaptic density because nearly every dopamine varicosity in the prefrontal cortex forms a synapse (Séguéla et al., 1988). Varicosities also represent sites where neurotransmitter synthesis, packaging, release, and reuptake most often occur (Benes et al., 1996). Stereology was conducted as previously described (Manitt et al., 2011; Reynolds et al., 2022). Contours of the dense TH-positive innervation in the medial prefrontal cortex were traced at x5 magnification using a Leica DM400B microscope and StereoInvestigator software. To determine the number of TH-positive varicosities, we used the optical fractionator probe function of Stereoinvestigator with a grid of 175 $um^2$, a counting frame of 25 $um^2$, a counting depth of 10 um, and a guard zone of 5 um. Counting was conducted at x100 magnification using the standard counting protocol for quantifying discrete objects (“particle stereology”) (Howard and Reed, 2004). Counts were performed blind by one individual per experiment (DH, AH, TO, or AD depending on the experiment). To determine the volume of each subregion we used the Cavalieri method in Stereoinvestigator (Howard and Reed, 2004). The coefficient of error was below 0.1 for all measures. Varicosity density was determined by dividing the total count of varicosities by the volume of the subregion.

**Stereotaxic Surgery**

To experimentally knock-down Netrin-1 along the dopamine axon route from the nucleus accumbens to the medial prefrontal cortex, we injected a Netrin-1 shRNA-expressing lentivirus or a scrambled control virus into the dorsal peduncular cortex.

Pre-designed and validated siRNA sequences (Ambion) were used to create shRNA (sequence GGAGCUCUAUAAGCUAUCA) by the addition of a standard hairpin loop (TTCAAGAGA) between the sense and antisense sequences. A scrambled control was created by rearranging the sequence order so that there was less than a 64% interaction rate. Active or control shRNA sequences were cloned into a pLentiLox 3.7 vector (Addgene, Plasmid #11795). Lentiviruses expressing the shRNAs and scrambled controls were prepared by the SPARC Biocentre lentiviral core facility (SickKids Hospital, Toronto, ON, Canada). For more details and validation information, see (Cuesta et al., 2020).

21-day-old mice were anesthetized with isoflurane (5% for induction and 2% for maintenance) and placed in a stereotaxic apparatus. Using Hamilton syringes, the shRNA-expressing lentivirus, or the lentivirus expressing the scrambled control sequence, were microinfused bilaterally into the dorsal peduncular cortices stereotaxically using the coordinates: +2.00 mm anterior/posterior, -0.05 mm medial/lateral, and −3.45 mm dorsal/ventral relative to Bregma. A total of 0.5 µl of purified virus was delivered on each side at an injection rate of 0.08 uL/min, which was then followed by a 3 min pause to allow of the virus to diffuse away from the syringe before the syringe was retracted. For anatomical experiments, the Netrin-1 knockdown and scrambled control viruses were injected into the left and right hemispheres, with the type of virus injected into each hemisphere determined randomly. For behavioural experiments, the same virus was injected into both hemispheres.

**Behaviour – *Go/No-Go***

We used the Go/No-Go task to measure inhibitory control, as we have described previously (Cuesta et al., 2019; Reynolds et al., 2018a, 2018b). The mice used for this experiment were adults (75 ± 15 days old at the beginning of the experiment) which had been stereotaxically injected with a Netrin-1 inhibiting or control virus at the onset of adolescence (see previous section).

During the experiment, mice were food restricted to 1.5 g food per to maintain a body weight of 85% of their initial free feeding weight. We used operant behavioral boxes (Med Associates, Inc., St Albans, VT, USA) equipped with a house light, an Sonalert tone generator, two illuminated nose poke holes, and a pellet dispenser. Chocolate-flavored dustless precision food pellets (BioServ, Inc., Flemington, NJ, USA) were used as our operant reinforcer. The experimental procedure consisted of two training stages, Discrimination Training and Reaction Time, followed by the Go/No-Go test phase. One session was conducted per mouse per day.

The first training stage is Discrimination Training. For this stage, at the start of each 20 min session, the house light comes on and remains illuminated throughout the session. Trials consist of the illumination of one nose poke hole for 9 seconds, counterbalanced between nose-poke holes across mice. If the mouse does not nose-poke into the illuminated hole within that 9 second period, the cue light is extinguished for a 10 second inter-trial interval before the next trial. If the mouse responds to the cue light by nose-poking, they received a pellet and the trial is counted as a “rewarded” trial. Responses to the active nose poke hole when the cue light is off, as well as responses to the non-active nose poke hole (where the cue light was never illuminated), were not rewarded. Mice received one Discrimination Training  session per day until they reached a rate of 70% rewarded trials, at which point they advanced to the next stage of training.

The second training stage is Reaction Time. At this stage, mice were trained to respond only within 3 seconds of the cue illumination to receive the pellet reward. These training sessions lasted 30 min, but the house light does not remain illuminated throughout the session. Instead, the house light becomes illuminated for a variable amount of time (3, 6, or 9 seconds, distributed randomly) prior to the illumination of the cue light, to signal the start of a new trial. This is designed to signal for the mice to attend to the cue. If the mice responded during this pretrial period (a ‘Premature Response’), the house light was turned off for a 10 second inter-trial interval and then a new trial is initiated. If the mouse did not perform a Premature Response, the cue light was illuminated for 3 seconds. A nose poke into the illuminated hole during this 3 second period resulted in the delivery of a reward pellet. If the mouse did not respond, the cue and house lights were extinguished and a 10 second inter-trial interval was initiated, followed by a new trial. Mice received one Reaction Time training session per day until they reached a rate of 70% rewarded trials and fewer than 25% of trials ended due to a Premature Response, at which point they advanced to the Go/No-Go test stage.

After training, mice underwent 10 daily sessions of the Go/No-Go task. This task required the mice to respond to the illuminated cue light (a “Go” trial) or to inhibit their response to this cue when it was presented in tandem with an 80 dB tone (a “No-Go” trial) to receive a reward pellet. During a “No-Go” trial, if mice responded during the 3 second presentation of both the illumination and tone cues, a 10 second inter-trial interval was initiated, followed by a new trial. A randomized, variable period of 3–9 seconds during which only the house light was illuminated signaled the start of each trial. A nose-poke during this time initiated a 10 second inter-trial period followed by a new trial. Within each session, the number of “Go” and “No-Go” trials were given in an approximately 1:1 ratio and presented in a randomized order. Each session lasted 30 min and consisted of approximately 60-100 completed trials.

We quantified three measures from the Go/No-Go Test data. Commission errors were our measure of inhibitory control. A commission error occurs when a mouse nose-pokes during a “No-Go” trial, when the cue light is illuminated concurrently with the 80 dB tone. We also quantified omission errors, which are when a mouse fails to nose-poke during a “Go” trial, when the cue light is illuminated in the absence of the tone. Finally, we calculated the correct response rate, which is the number of “Go” trials where the mouse nose-pokes while the cue light is illuminated plus the number of “No-Go” trials where the mouse does not nose-poke while the cue light is illuminated. All three measures are analysed as proportions of the total number of trials presented each test day.

**Behaviour – *Light/Dark Box***

In hamsters, we used the light/dark box test, as we’ve described previously (Kyne et al., 2019). We used operant behavioural boxes consisting of two compartments: one with illumination from a house light (the light compartment; 40.0 cm × 39.9 cm × 31.2 cm) and one without illumination (the dark compartment; 38.9 cm × 12.7 cm × 15.2 cm). The compartments were separated by barrier with an opening that could be blocked by a metal door.

For each session, a hamster placed inside the dark compartment of the apparatus with the metal door closed. The session was initiated when the metal door was opened, allowing the hamster to explore the light compartment. The hamster was allowed to move freely between the two compartments for 10 minutes. We used the amount of time spent in the light compartment as our measure of exploratory behaviour.

The hamsters were recorded by a camera mounted above the boxes using Media Recorder 4 software (Noldus Information Technology Inc., Wageningen, The Netherlands). Scoring was done automatically using EthoVision XT10 software (Noldus Information Technology Inc., Wageningen, The Netherlands).

**Statistical Analyses**

Detailed statistical explanations for each analysis are presented in our Statistics Supplement. All analyses were conducted in the statistical programming language R (Team, 2014). For all analyses our significance threshold was set at p = 0.05. All code and data necessary to reproduce the statistical analyses presented in our manuscript are available for download from the Open Science Framework (DIO 10.17605/OSF.IO/DU3H4).
