## Supplementary Statistics for "The scheduling of adolescence with Netrin-1 and UNC5C"

### Statistics Supplement

Daniel Hoops

#### Figure 1, panel D

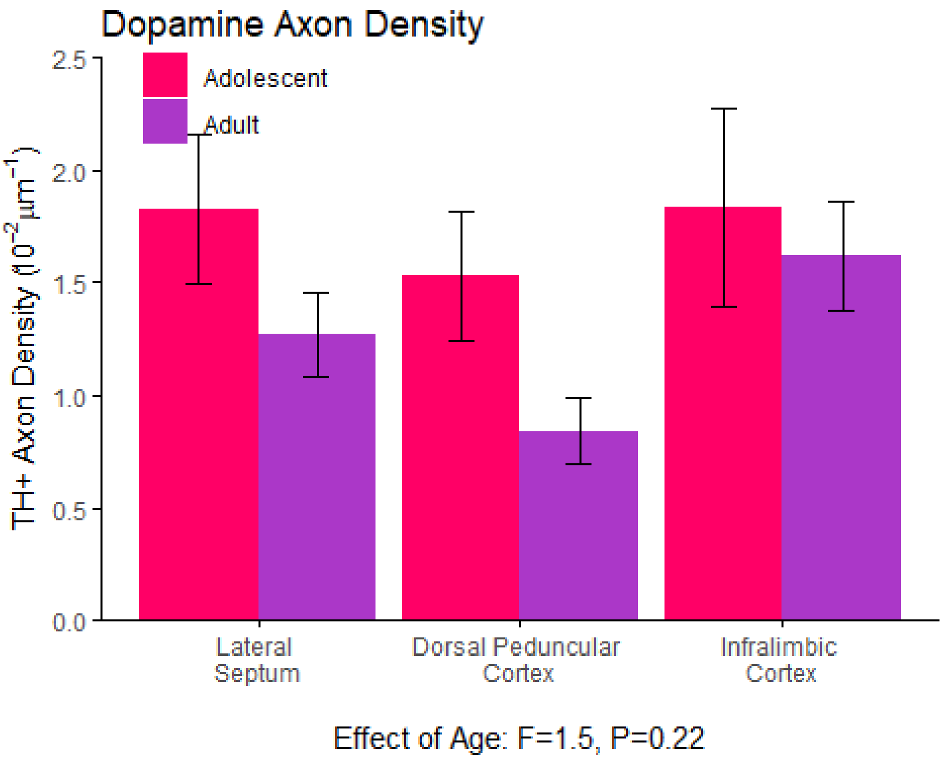


This panel illustrates the results of the quantification of dopamine axons along the dopamine axon bundle that runs from the nucleus accumbens to the medial prefrontal cortex, through the septum and adjacent structures. The modified stereological approach we used is described in detail in our methods supplement and by Kim et al., 2011. It is an unbiased method for counting elongated structures such as axons.

*Summary Statistics:*

| age | region | N | density.graph | sd | se | ci |
|---|---|---|---|---|---|---|
| Adolescent | Dorsal Peduncular Cortex | 9 | 1.53 | 0.86 | 0.29 | 0.66 |
| Adolescent | Infralimbic Cortex | 10 | 1.84 | 1.39 | 0.44 | 0.99 |
| Adolescent | Lateral Septum | 11 | 1.83 | 1.09 | 0.33 | 0.74 |
| Adult | Dorsal Peduncular Cortex | 9 | 0.84 | 0.44 | 0.15 | 0.34 |
| Adult | Infralimbic Cortex | 9 | 1.62 | 0.72 | 0.24 | 0.56 |
| Adult | Lateral Septum | 9 | 1.27 | 0.56 | 0.19 | 0.43 |

Significant differences in the density of the dopamine axon bundle between adolescence (21 days old) and adulthood (75 days old) were detected using an analysis of variance (ANOVA). We calculated a mixed-effects ANOVA with brain region, hemisphere, and age as fixed effects, a region-by-age interaction, mouse ID as a random effect, and axon density as the response variable.

```
## Analysis of Deviance Table (Type II tests)
##
## Response: scale.density
##              Chisq Df Pr(>Chisq)
## hemisphere 27.4459  1  1.615e-07 ***
## region      6.2703  2    0.04349 *
## age         1.5368  1    0.21510
## region:age  1.4359  2    0.48776
## ---
## Signif. codes:  0 '***' 0.001 '**' 0.01 '*' 0.05 '.' 0.1 ' ' 1
```

#### Figure 1, panel E

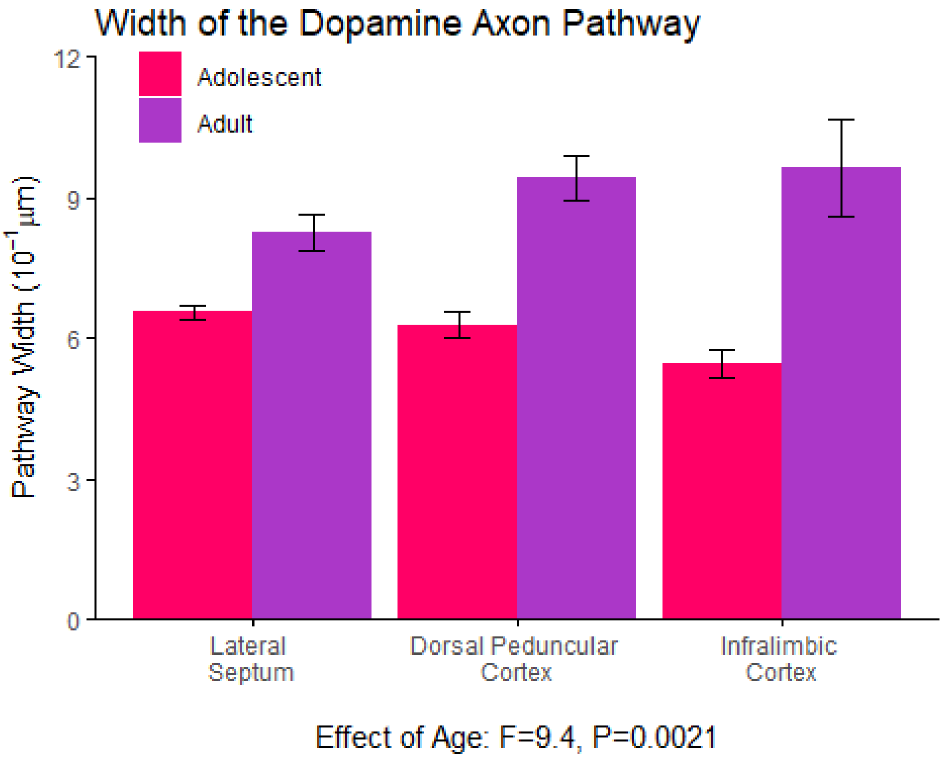


This panel illustrates the difference in the width of the dopamine axon bundle between adolescence and adulthood. This data originates from the same samples as the data in panel 1D.

*Summary Statistics:*

| age | region | N | width.graph | sd | se | ci |
|---|---|---|---|---|---|---|
| Adolescent | Dorsal Peduncular Cortex | 9 | 6.28 | 0.86 | 0.29 | 0.66 |
| Adolescent | Infralimbic Cortex | 10 | 5.45 | 0.94 | 0.30 | 0.67 |
| Adolescent | Lateral Septum | 11 | 6.55 | 0.47 | 0.14 | 0.31 |
| Adult | Dorsal Peduncular Cortex | 9 | 9.40 | 1.45 | 0.48 | 1.11 |
| Adult | Infralimbic Cortex | 9 | 9.64 | 3.13 | 1.04 | 2.40 |
| Adult | Lateral Septum | 9 | 8.25 | 1.15 | 0.38 | 0.89 |

Significant differences in the width of the dopamine axon bundle between adolescence (21 days old) and adulthood (75 days old) were detected using an analysis of variance (ANOVA). We calculated a mixed-effects ANOVA with brain region, hemisphere, and age as fixed effects, a region-by-age interaction, mouse ID as a random effect, and bundle width as the response variable.

```
## Analysis of Deviance Table (Type II tests)
##
```

```
## Response: scale.width
##             Chisq Df Pr(>Chisq)
## hemisphere 0.1443  1   0.704081
## region     1.3000  2   0.522055
## age        9.7304  1   0.001812 **
## region:age 5.7381  2   0.056752 .
## ---
## Signif. codes:  0 '***' 0.001 '**' 0.01 '*' 0.05 '.' 0.1 ' ' 1
```

#### Figure 1, panel H

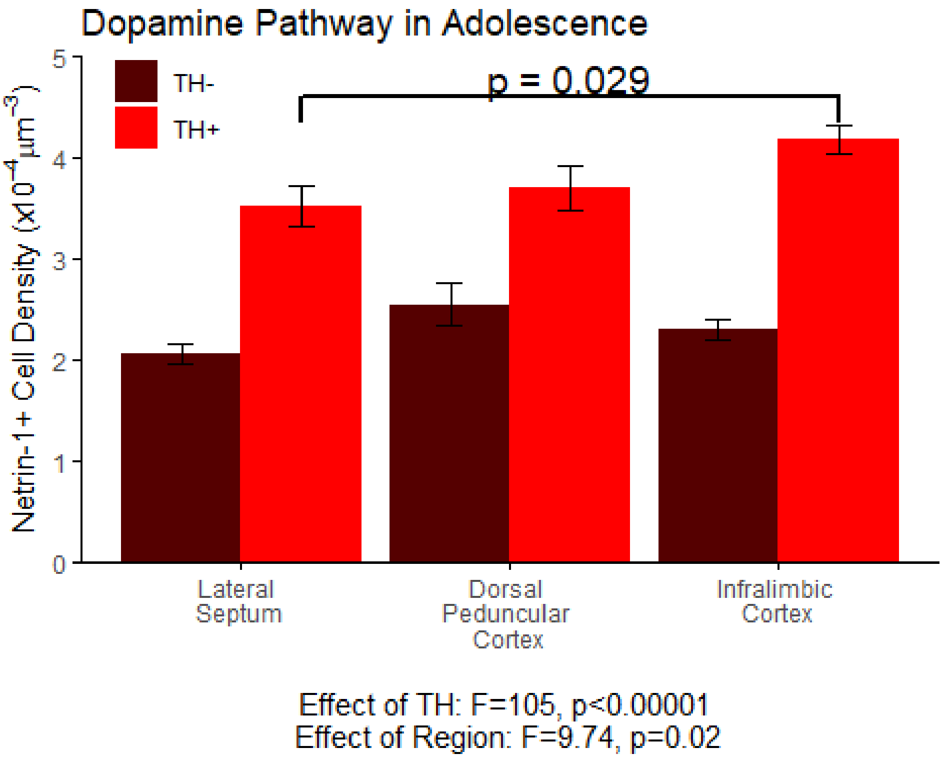


This panel illustrates the results of the quantification of Netrin-1 positive cells along the dopamine axon bundle ("TH+") and in an area of equal size immediately medial to it ("TH-"). The density of Netrin-1 positive cells in each area was quantified using standard unbiased stereology. This analysis was conducted in early adolescent (21-day-old) mice.

*Summary Statistics:*

| TH | Region | N | Density.graph | sd | se | ci |
|---|---|---|---|---|---|---|
| TH- | Dorsal Peduncular Cortex | 8 | 2.54 | 0.59 | 0.21 | 0.50 |
| TH- | Infralimbic Cortex | 8 | 2.30 | 0.28 | 0.10 | 0.23 |
| TH- | Lateral Septum | 8 | 2.07 | 0.28 | 0.10 | 0.24 |
| TH+ | Dorsal Peduncular Cortex | 8 | 3.70 | 0.63 | 0.22 | 0.52 |
| TH+ | Infralimbic Cortex | 8 | 4.18 | 0.39 | 0.14 | 0.33 |
| TH+ | Lateral Septum | 8 | 3.51 | 0.57 | 0.20 | 0.47 |

Significant differences in the density of the Netrin-1 positive cells were detected using an analysis of variance (ANOVA). We calculated a mixed-effects ANOVA with brain region, hemisphere, and TH as fixed effects, mouse ID as a random effect, and Netrin-1 positive cell density as the response variable. We detected a main effect of TH (whether the area overlapped with the dopamine axon bundle) as well as a main effect of region.

```
## Analysis of Deviance Table (Type II tests)
##
## Response: Density
##               Chisq Df Pr(>Chisq)
## Hemisphere   0.8838  1    0.34716
## TH         106.6570  1    < 2e-16 ***
## Region       6.8505  2    0.03254 *
## ---
## Signif. codes:  0 '***' 0.001 '**' 0.01 '*' 0.05 '.' 0.1 ' ' 1
```

Because in this analysis there are significant effects of region and TH, we conducted post-hoc Tukey tests separately for the TH-positive and TH-negative data to determine which regions have different densities of Netrin-1-positive cells. We found that the only difference in density occurs between the lateral septum and infralimbic cortex along the dopamine axon route. This indicates that, overlapping with the dopamine axon bundle, there is an increasing density of Netrin-1 positive cells towards the dopamine innervation target (the medial prefrontal cortex).

```
## $`emmeans of Region`
##   Region                          emmean      SE df lower.CL upper.CL
##  Lateral \nSeptum             0.000351 4.2e-05  3 0.000218 0.000485
##  Dorsal \nPeduncular \nCortex 0.000370 4.2e-05  3 0.000236 0.000504
##  Infralimbic \nCortex         0.000418 4.2e-05  3 0.000284 0.000551
##
## Results are averaged over the levels of: Hemisphere
## Degrees-of-freedom method: containment
## Confidence level used: 0.95
##
## $`pairwise differences of Region`
##     1                                                 estimate       SE df
##  Lateral \nSeptum - Dorsal \nPeduncular \nCortex     -1.86e-05 2.33e-05 17
##  Lateral \nSeptum - Infralimbic \nCortex             -6.63e-05 2.33e-05 17
##  Dorsal \nPeduncular \nCortex - Infralimbic \nCortex -4.78e-05 2.33e-05 17
##  t.ratio p.value
##   -0.796  0.7106
##   -2.845  0.0286
##   -2.049  0.1308
##
## Results are averaged over the levels of: Hemisphere
## Degrees-of-freedom method: containment
## P value adjustment: tukey method for comparing a family of 3 estimates
```

```
## $`emmeans of Region`
##   Region                          emmean       SE df lower.CL upper.CL
##  Lateral \nSeptum             0.000207 2.38e-05  3 0.000131 0.000282
##  Dorsal \nPeduncular \nCortex 0.000254 2.38e-05  3 0.000179 0.000330
##  Infralimbic \nCortex         0.000230 2.38e-05  3 0.000154 0.000305
##
## Results are averaged over the levels of: Hemisphere
## Degrees-of-freedom method: containment
```

```
## Confidence level used: 0.95
##
## $`pairwise differences of Region`
##     1                                                  estimate       SE df
##  Lateral \nSeptum - Dorsal \nPeduncular \nCortex     -4.78e-05 1.86e-05 17
##  Lateral \nSeptum - Infralimbic \nCortex             -2.30e-05 1.86e-05 17
##  Dorsal \nPeduncular \nCortex - Infralimbic \nCortex  2.48e-05 1.86e-05 17
##  t.ratio p.value
##   -2.564  0.0501
##   -1.236  0.4491
##    1.328  0.3995
##
## Results are averaged over the levels of: Hemisphere
## Degrees-of-freedom method: containment
## P value adjustment: tukey method for comparing a family of 3 estimates
```

#### Figure 1, panel I

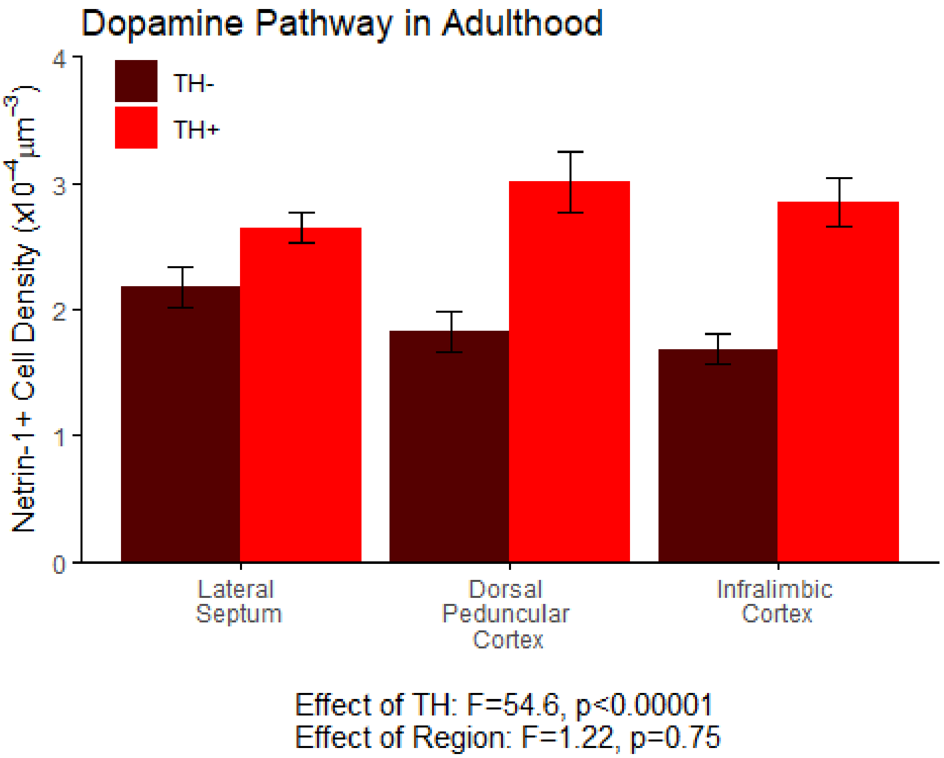


This panel illustrates the results of the quantification of Netrin-1 positive cells along the dopamine axon bundle ("TH+") and in an area of equal size immediately medial to it ("TH-"). The density of Netrin-1 positive cells in each area was quantified using standard unbiased stereology. This analysis was conducted in adult (75 day old) mice.

*Summary Statistics:*

| TH | Region | N | Density.graph | sd | se | ci |
|---|---|---|---|---|---|---|
| TH- | Dorsal Peduncular Cortex | 8 | 1.82 | 0.44 | 0.16 | 0.37 |
| TH- | Infralimbic Cortex | 8 | 1.69 | 0.35 | 0.12 | 0.29 |
| TH- | Lateral Septum | 8 | 2.18 | 0.45 | 0.16 | 0.38 |
| TH+ | Dorsal Peduncular Cortex | 8 | 3.01 | 0.69 | 0.24 | 0.58 |
| TH+ | Infralimbic Cortex | 8 | 2.84 | 0.54 | 0.19 | 0.45 |
| TH+ | Lateral Septum | 8 | 2.65 | 0.32 | 0.11 | 0.27 |

Significant differences in the density of the Netrin-1 positive cells were not detected in adulthood using an analysis of variance (ANOVA). We calculated a mixed-effects ANOVA with brain region, hemisphere, and TH as fixed effects, mouse ID as a random effect, and Netrin-1 positive cell density as the response variable.

```
## Analysis of Deviance Table (Type II tests)
##
```

```
## Response: Density
##               Chisq Df Pr(>Chisq)    
## Hemisphere  3.0255  1    0.08196 .  
## TH         52.8899  1  3.528e-13 ***
## Region      1.1838  2    0.55327    
## ---
## Signif. codes:  0 '***' 0.001 '**' 0.01 '*' 0.05 '.' 0.1 ' ' 1
```

#### Figure 1, panel L

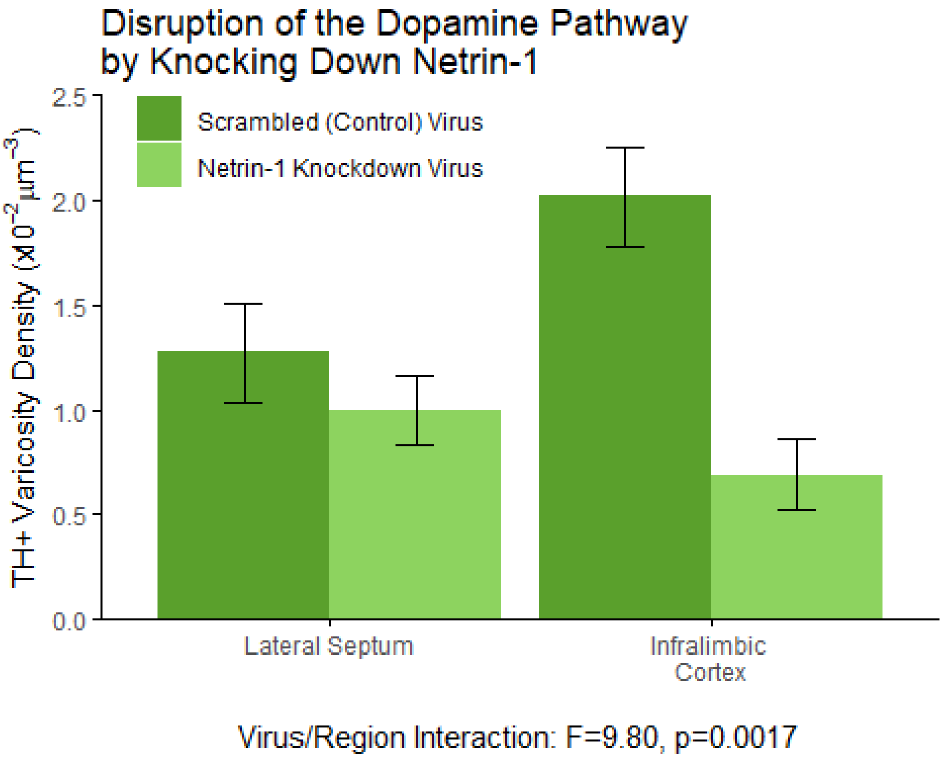


This panel illustrates the results of the quantification of TH-positive varicosities along the route by which dopamine axons grow to the medial prefrontal cortex. At the onset of adolescence (21 days old), these mice were injected with either a virus to knock down Netrin-1 or a nonfunctional control virus into the dorsal peduncular cortex, a structure midway along to the dopamine axon route. Each mouse received the knock-down virus in one hemisphere and the control virus in the other hemisphere. Virus assignment to each hemishere was random.

Once the mice reached adulthood, 75 days old, we quantified TH-positive varicosities in the lateral septum, a region growing dopamine axons reach before the location of the injection, and in the infralimbic cortex, a region the growing dopamine axons reach after the location of the injection. The density of TH-positive varicosities in each area was quantified using standard unbiased stereology.

*Summary Statistics:*

| virus | region | N | density.graph | sd | se | ci |
|---|---|---|---|---|---|---|
| Netrin-1 Knockdown Virus | Infralimbic Cortex | 11 | 0.69 | 0.55 | 0.17 | 0.37 |
| Netrin-1 Knockdown Virus | Lateral Septum | 11 | 1.00 | 0.54 | 0.16 | 0.37 |
| Scrambled (Control) Virus | Infralimbic Cortex | 8 | 2.01 | 0.68 | 0.24 | 0.57 |
| Scrambled (Control) Virus | Lateral Septum | 8 | 1.27 | 0.68 | 0.24 | 0.57 |

Significant differences in the density of the TH-positive varicosities were detected using an analysis of variance (ANOVA). We calculated a mixed-effects ANOVA with brain region, hemisphere, and TH as fixed effects, mouse ID as a random effect, and Netrin-1 positive cell density as the response variable.

```
## Analysis of Deviance Table (Type II tests)
##
## Response: scale.density
##                Chisq Df Pr(>Chisq)
## hemisphere    1.9885  1  0.1584929
## sex           1.3527  1  0.2448092
## region        0.6696  1  0.4131824
## virus        13.1505  1  0.0002874 ***
## region:virus  9.8023  1  0.0017429 **
## ---
## Signif. codes:  0 '***' 0.001 '**' 0.01 '*' 0.05 '.' 0.1 ' ' 1
```

Post-hoc tukey tests reveal that the the density of TH-positive varicosities in the infralimbic cortex is lower where the Netrin-1 knockdown virus was injected compared to where the control virus was injected. No effect is observed in the lateral septum.

Lateral Septum:

```
## $`emmeans of virus`
##  virus                     emmean    SE df lower.CL upper.CL
##  Scrambled (Control) Virus  0.176 0.286  2    -1.05    1.404
##  Netrin-1 Knockdown Virus  -0.159 0.269  2    -1.31    0.997
##
## Results are averaged over the levels of: hemisphere
## Degrees-of-freedom method: containment
## Confidence level used: 0.95
##
## $`pairwise differences of virus`
##  1                                                  estimate    SE df
##  Scrambled (Control) Virus - (Netrin-1 Knockdown Virus)    0.335 0.363  2
##  t.ratio p.value
##    0.923  0.4535
##
## Results are averaged over the levels of: hemisphere
## Degrees-of-freedom method: containment
```

Infralimbic Cortex:

```
## $`emmeans of virus`
##  virus                     emmean    SE df lower.CL upper.CL
##  Scrambled (Control) Virus  1.108 0.217  2    0.175    2.041
##  Netrin-1 Knockdown Virus  -0.607 0.205  2   -1.491    0.276
##
## Results are averaged over the levels of: hemisphere
## Degrees-of-freedom method: containment
## Confidence level used: 0.95
```

```
##
## $`pairwise differences of virus`
##  1                                                      estimate    SE df
##  Scrambled (Control) Virus - (Netrin-1 Knockdown Virus)     1.72 0.269  2
##  t.ratio p.value
##    6.376  0.0237
##
## Results are averaged over the levels of: hemisphere
## Degrees-of-freedom method: containment
```

#### Figure 1, panel N

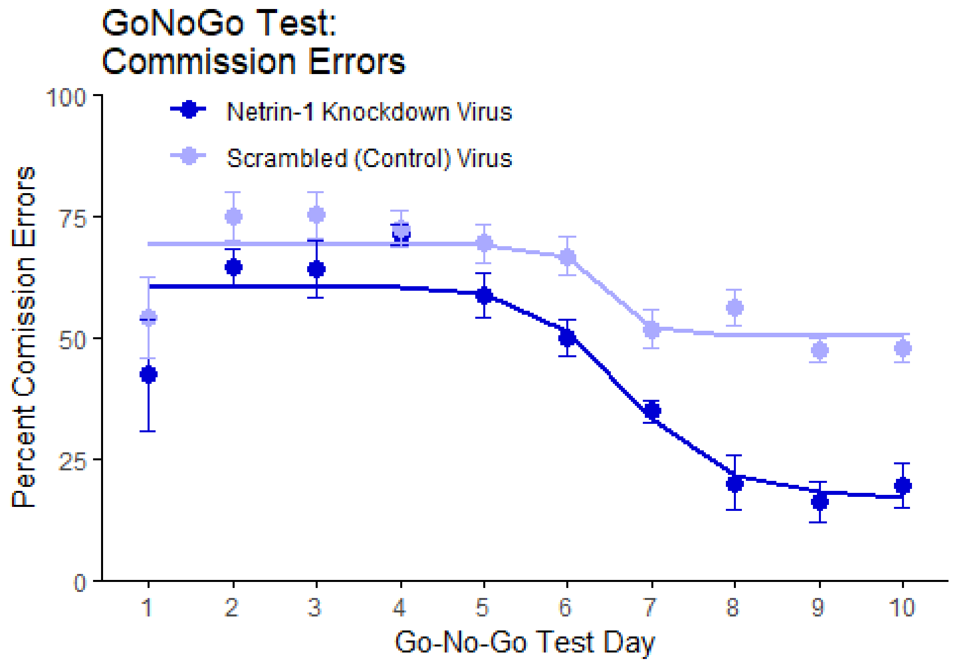


This panel illustrates the ability of adult (at least 60-day-old) mice to withhold their response to a stimulus in order to receive a food reward during the GoNoGo behavioural test. Inhibitory control is quantified as the percentage of trials during which the mice should have witheld their response, but failed to ("commission errors").

At the onset of adolescence (21 days old) the mice used in this experiment received bilateral injections of a virus that either knocked down Netrin-1 or served as a nonfunctional control. The viruses were injected along the route by which dopamine axons grow from the nucleus accumbens to the medial prefrontal cortex during adolescence (see methods for details).

*Summary Statistics:*

| virus | Day | N | Percent.Comission.Errors | sd | se | ci |
|---|---|---|---|---|---|---|
| Netrin-1 Knockdown Virus | 1 | 10 | 0.42 | 0.36 | 0.11 | 0.26 |
| Netrin-1 Knockdown Virus | 2 | 10 | 0.64 | 0.13 | 0.04 | 0.09 |
| Netrin-1 Knockdown Virus | 3 | 10 | 0.64 | 0.18 | 0.06 | 0.13 |
| Netrin-1 Knockdown Virus | 4 | 10 | 0.71 | 0.06 | 0.02 | 0.05 |
| Netrin-1 Knockdown Virus | 5 | 10 | 0.59 | 0.14 | 0.05 | 0.10 |
| Netrin-1 Knockdown Virus | 6 | 10 | 0.50 | 0.12 | 0.04 | 0.08 |
| Netrin-1 Knockdown Virus | 7 | 10 | 0.35 | 0.07 | 0.02 | 0.05 |

| virus | Day | N | Percent.Comission.Errors | sd | se | ci |
|---|---|---|---|---|---|---|
| Netrin-1 Knockdown Virus | 8 | 10 | 0.20 | 0.18 | 0.06 | 0.13 |
| Netrin-1 Knockdown Virus | 9 | 10 | 0.16 | 0.13 | 0.04 | 0.09 |
| Netrin-1 Knockdown Virus | 10 | 8 | 0.20 | 0.13 | 0.05 | 0.11 |
| Scrambled (Control) Virus | 1 | 10 | 0.54 | 0.26 | 0.08 | 0.19 |
| Scrambled (Control) Virus | 2 | 10 | 0.75 | 0.16 | 0.05 | 0.12 |
| Scrambled (Control) Virus | 3 | 10 | 0.75 | 0.15 | 0.05 | 0.11 |
| Scrambled (Control) Virus | 4 | 10 | 0.73 | 0.12 | 0.04 | 0.08 |
| Scrambled (Control) Virus | 5 | 10 | 0.70 | 0.13 | 0.04 | 0.09 |
| Scrambled (Control) Virus | 6 | 10 | 0.67 | 0.12 | 0.04 | 0.09 |
| Scrambled (Control) Virus | 7 | 10 | 0.52 | 0.13 | 0.04 | 0.09 |
| Scrambled (Control) Virus | 8 | 10 | 0.56 | 0.12 | 0.04 | 0.09 |
| Scrambled (Control) Virus | 9 | 10 | 0.48 | 0.08 | 0.03 | 0.06 |
| Scrambled (Control) Virus | 10 | 10 | 0.48 | 0.09 | 0.03 | 0.07 |

An analysis of variance (ANOVA) revealed a significant difference between mice that were bilaterally injected with the Netrin-1 knockdown virus and those that were injected with the nonfunctional control virus. We determined this using a mixed-effects ANOVA with virus and day as fixed effects, mouse ID as a random effect, and percent of responses that were commission errors as the response variable.

```
## Analysis of Deviance Table (Type II tests)
##
## Response: Percent.Comission.Errors
##             Chisq Df Pr(>Chisq)
## Day       68.3153  1  < 2.2e-16 ***
## virus      3.0905  1   0.078751 .
## Day:virus  9.0026  1   0.002696 **
## ---
## Signif. codes:  0 '***' 0.001 '**' 0.01 '*' 0.05 '.' 0.1 ' ' 1
```

#### Figure 1, panel O

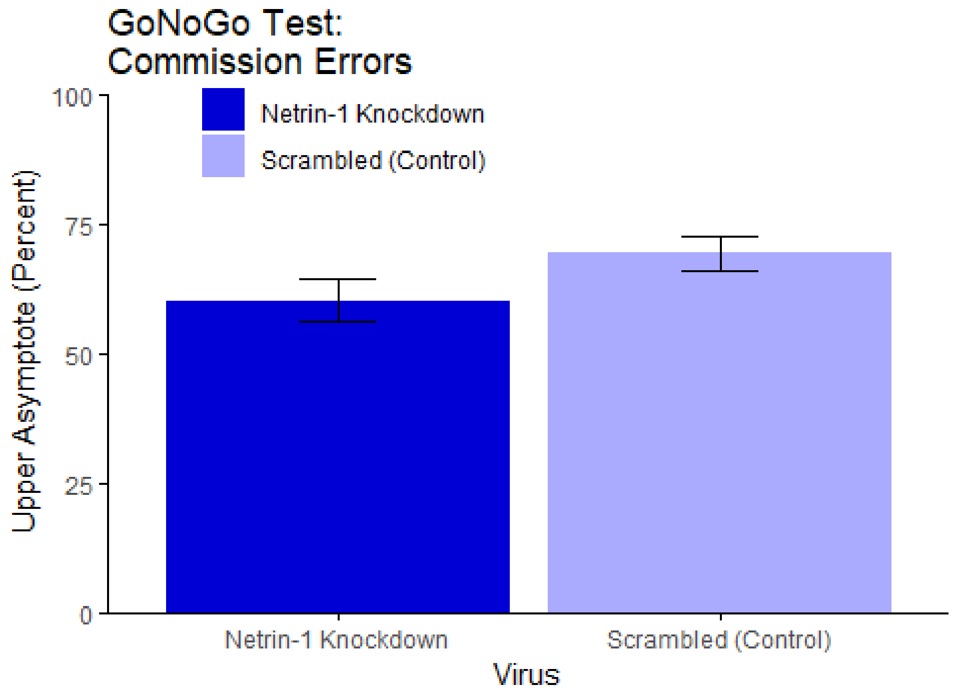


t=5.18, p<0.0001

This panel illustrates the rate at which mice fail to withhold their response to a stimulus during the initial phase of the GoNoGo Test.

*Summary Statistics:*

| component | virus | Estimate | sd |
|---|---|---|---|
| upper asymptote | Netrin-1 Knockdown | 0.60 | 0.04 |
| upper asymptote | Scrambled (Control) | 0.69 | 0.03 |

A t-test revealed a significant difference between viral treatments in the rate of commission errors during the initial part of the GoNoGo test. We determined this using a two-way t-test with viral treatment (Netrin-1 knockdown or nonfunctional control virus) as the independent variable and percent of responses that were commission errors as the response variable.

```
## Difference of means             Std Error              t-value             p-
value
##       -9.003540e-02        1.737648e-02        -5.181451e+00           7.17098
9e-05
```

#### Figure 1, panel P

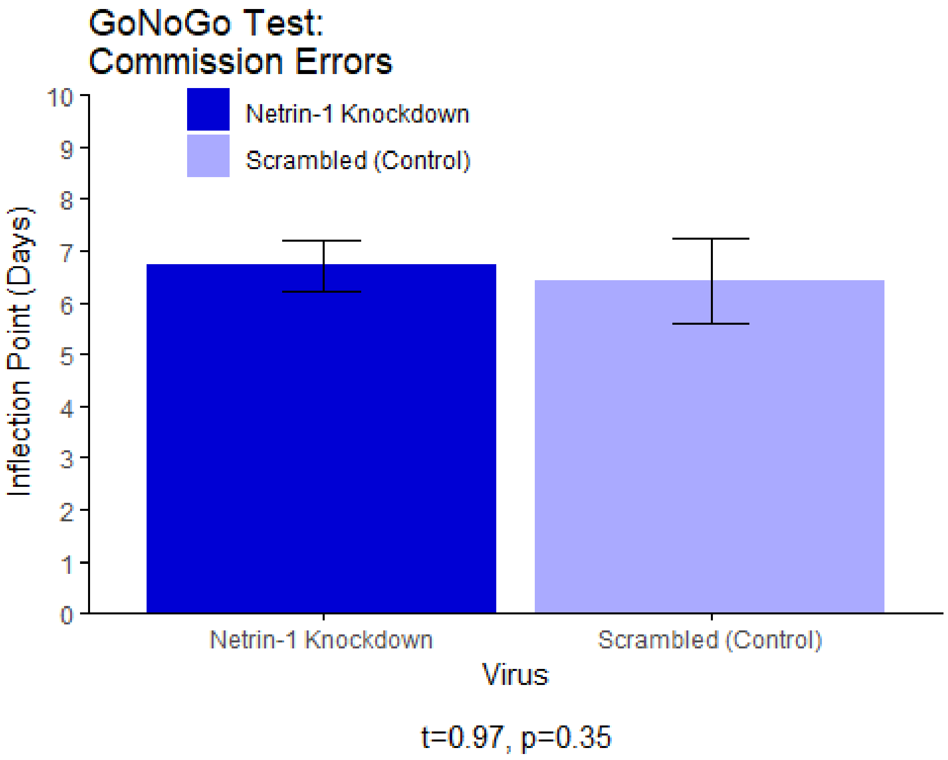


This panel illustrates the number of days to the inflection point (ED50) of the sigmoidal curves shown in panel 1N.

*Summary Statistics:*

| component | virus | Estimate | sd |
|---|---|---|---|
| ED50 | Netrin-1 Knockdown | 6.71 | 0.50 |
| ED50 | Scrambled (Control) | 6.42 | 0.81 |

A t-test revealed no significant difference between viral treatments in the time to the inflection point of the GoNoGo test. We determined this using two-way t-test with virus as the independent variable and time (in days) of the inflection point as the response variable.

```
## Difference of means           Std Error              t-value               p-
value
##          0.2931987           0.3009151            0.9743568             0.34
53438
```

#### Figure 1, panel Q

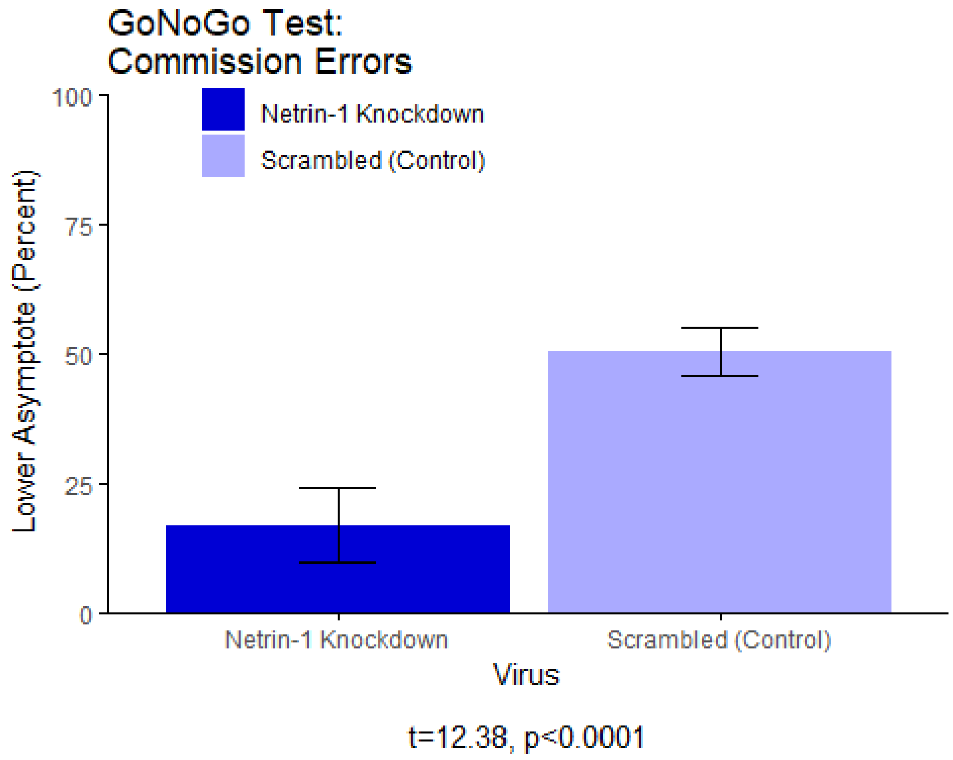


This panel illustrates the rate at which mice fail to withhold their response to a stimulus during the final phase of the GoNoGo Test.

*Summary Statistics:*

| component | virus | Estimate | sd |
|---|---|---|---|
| lower asymptote | Netrin-1 Knockdown | 0.17 | 0.07 |
| lower asymptote | Scrambled (Control) | 0.50 | 0.05 |

A t-test revealed a significant difference between viral treatments in the rate of commission errors during the final part of the GoNoGo test. We determined this using two-way t-test with viral treatment as the independent variable and percent of responses that were commission errors as the response variable.

```
## Difference of means          Std Error           t-value             p-
value
##       -3.347033e-01       2.703853e-02       -1.237875e+01          1.63080
7e-09
```

#### Figure 3, panel C

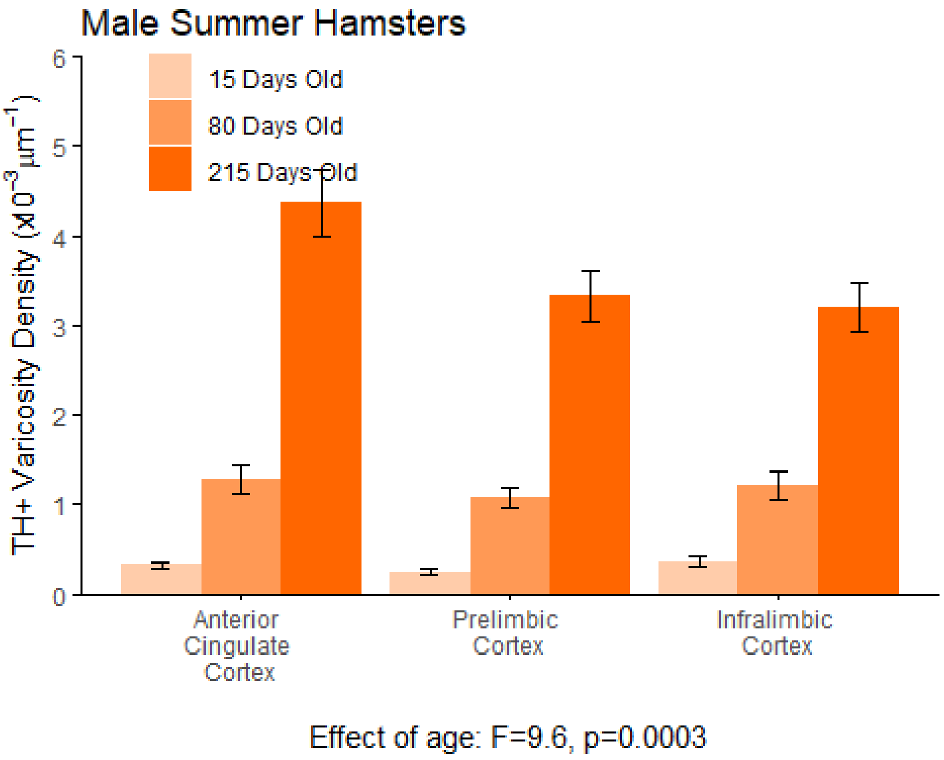


This panel illustrates the results of the quantification of the density of TH-positive varicosities in the dopamine-innervated part of the medial prefrontal cortex. This analysis was conducted in 15-, 80-, and 215-day-old male hamsters that were housed under a summer-mimicking light cycle. The density of dopamine varicosities in each area was quantified using standard unbiased stereology.

*Summary Statistics:*

| age | region | N | density.graph | sd | se | ci |
|---|---|---|---|---|---|---|
| 15 Days Old | Anterior Cingulate Cortex | 8 | 0.32 | 0.09 | 0.03 | 0.07 |
| 15 Days Old | Infralimbic Cortex | 8 | 0.36 | 0.15 | 0.05 | 0.13 |
| 15 Days Old | Prelimbic Cortex | 8 | 0.25 | 0.11 | 0.04 | 0.09 |
| 215 Days Old | Anterior Cingulate Cortex | 10 | 4.37 | 1.18 | 0.37 | 0.85 |
| 215 Days Old | Infralimbic Cortex | 10 | 3.21 | 0.85 | 0.27 | 0.61 |
| 215 Days Old | Prelimbic Cortex | 10 | 3.33 | 0.88 | 0.28 | 0.63 |
| 80 Days Old | Anterior Cingulate Cortex | 8 | 1.28 | 0.43 | 0.15 | 0.36 |
| 80 Days Old | Infralimbic Cortex | 8 | 1.21 | 0.45 | 0.16 | 0.38 |
| 80 Days Old | Prelimbic Cortex | 8 | 1.07 | 0.32 | 0.11 | 0.27 |

Using an analysis of variance (ANOVA), we found that the density of dopamine varicosities in the medial prefrontal cortex increases between age groups in summer male hamsters.

We calculated a mixed-effects ANOVA with brain region, age, and the time each sample was stored in the fridge before staining as fixed effects, hamster ID as a random effect, and varicosity density as the response variable.

```
##
## Error: hamster_ID
##     Df Sum Sq Mean Sq
## age  1  29.51   29.51
##
## Error: hamster_ID:region
##        Df Sum Sq Mean Sq
## region  2  0.431  0.2155
##
## Error: Within
##                              Df Sum Sq Mean Sq F value   Pr(>F)
## region                        2  1.141   0.571   2.350 0.104587
## age                           2  4.664   2.332   9.602 0.000255 ***
## fridge.scaled.age             1 22.528  22.528  92.767 1.46e-13 ***
## region:age                    4  0.178   0.045   0.184 0.946034
## region:fridge.scaled.age      2  1.000   0.500   2.058 0.137036
## age:fridge.scaled.age         2  3.588   1.794   7.388 0.001403 **
## region:age:fridge.scaled.age  4  0.114   0.029   0.118 0.975648
## Residuals                    57 13.842   0.243
## ---
## Signif. codes:  0 '***' 0.001 '**' 0.01 '*' 0.05 '.' 0.1 ' ' 1
```

Post-hoc tukey tests by age reveal that dopamine innervation to the medial prefrontal cortex increases between 15 and 80 days old, as well as between 80 and 215 days old.

```
##   Tukey multiple comparisons of means
##     95% family-wise confidence level
##
## Fit: aov(formula = scale.density ~ region * age * fridge.scaled.age, data
= MLD_data)
##
## $age
##                                diff        lwr       upr    p adj
## 80 Days Old-15 Days Old  0.3897021 0.03943209 0.7399721 0.025754
## 215 Days Old-15 Days Old 1.4854926 1.15319729 1.8177879 0.000000
## 215 Days Old-80 Days Old 1.0957905 0.76349522 1.4280858 0.000000
```

#### Figure 3, panel D

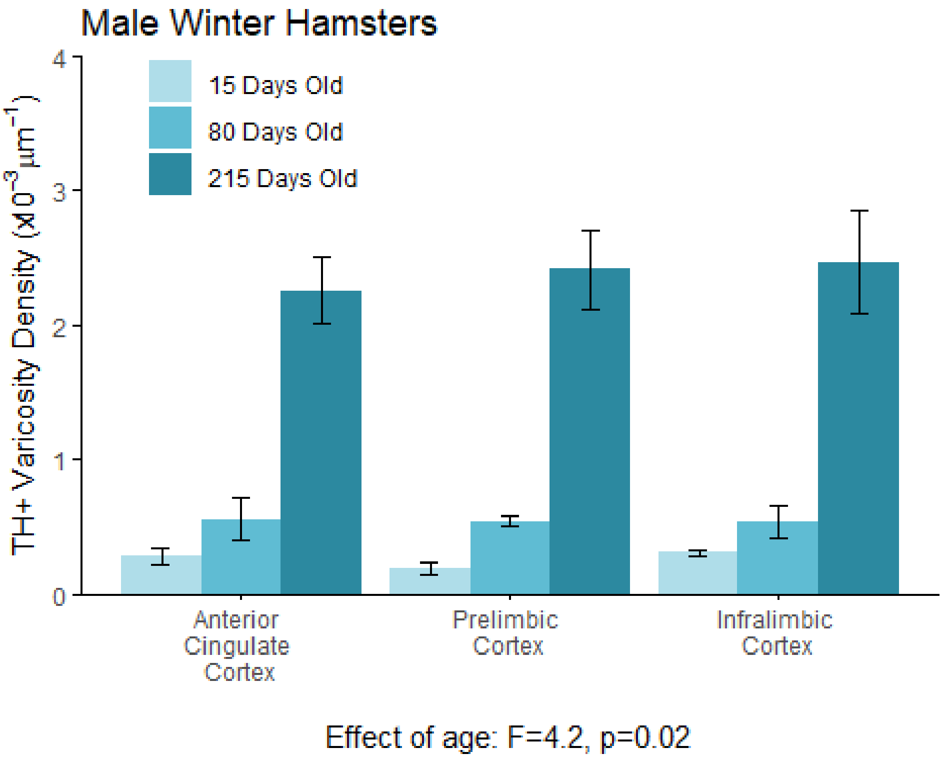


This panel illustrates the results of the quantification of the density of TH-positive varicosities in the dopamine-innervated part of the medial prefrontal cortex. This analysis was conducted in 15-, 80-, and 215-day-old male hamsters that were housed under a winter-mimicking light cycle. The density of dopamine varicosities in each area was quantified using standard unbiased stereology.

*Summary Statistics:*

| age | region | N | density.graph | sd | se | ci |
|---|---|---|---|---|---|---|
| 15 Days Old | Anterior Cingulate Cortex | 4 | 0.28 | 0.12 | 0.06 | 0.19 |
| 15 Days Old | Infralimbic Cortex | 4 | 0.30 | 0.04 | 0.02 | 0.06 |
| 15 Days Old | Prelimbic Cortex | 4 | 0.19 | 0.09 | 0.04 | 0.14 |
| 215 Days Old | Anterior Cingulate Cortex | 8 | 2.26 | 0.71 | 0.25 | 0.60 |
| 215 Days Old | Infralimbic Cortex | 8 | 2.47 | 1.09 | 0.39 | 0.91 |
| 215 Days Old | Prelimbic Cortex | 8 | 2.42 | 0.82 | 0.29 | 0.68 |
| 80 Days Old | Anterior Cingulate Cortex | 8 | 0.56 | 0.44 | 0.16 | 0.37 |
| 80 Days Old | Infralimbic Cortex | 8 | 0.54 | 0.34 | 0.12 | 0.28 |
| 80 Days Old | Prelimbic Cortex | 8 | 0.54 | 0.10 | 0.04 | 0.08 |

Using an analysis of variance (ANOVA), we found that the density of dopamine varicosities in the medial prefrontal cortex increases with age in winter male hamsters. We calculated a mixed-effects ANOVA with brain region, age, and the time each sample was stored in the fridge before staining as fixed effects, hamster ID as a random effect, and varicosity density as the response variable.

```
##
## Error: hamster_ID
##     Df Sum Sq Mean Sq
## age  1  16.78   16.78
##
## Error: hamster_ID:region
##        Df  Sum Sq Mean Sq
## region  2 0.02697 0.01349
##
## Error: Within
##                               Df Sum Sq Mean Sq F value  Pr(>F)
## region                         2   0.37   0.186   0.249 0.78067
## age                            2   6.25   3.123   4.167 0.02046 *
## fridge.scaled.age              1   0.02   0.018   0.025 0.87584
## region:age                     4   0.22   0.056   0.075 0.98955
## region:fridge.scaled.age       2   0.07   0.037   0.050 0.95154
## age:fridge.scaled.age          2  10.51   5.256   7.011 0.00189 **
## region:age:fridge.scaled.age   4   0.02   0.005   0.007 0.99990
## Residuals                     57  42.73   0.750
## ---
## Signif. codes:  0 '***' 0.001 '**' 0.01 '*' 0.05 '.' 0.1 ' ' 1
```

Post-hoc tukey tests by age reveal that dopamine innervation to the medial prefrontal cortex increases between 80 and 215 days old.

```
##   Tukey multiple comparisons of means
##     95% family-wise confidence level
##
## Fit: aov(formula = scale.density ~ region * age * fridge.scaled.age, data
= MSD_data)
##
## $age
##                                 diff        lwr      upr     p adj
## 80 Days Old-15 Days Old  -0.2448559 -0.8016999 0.311988 0.5444274
## 215 Days Old-15 Days Old  0.7147839  0.1579399 1.271628 0.0085256
## 215 Days Old-80 Days Old  0.9596399  0.3726748 1.546605 0.0006452
```

#### Figure 3, panel K

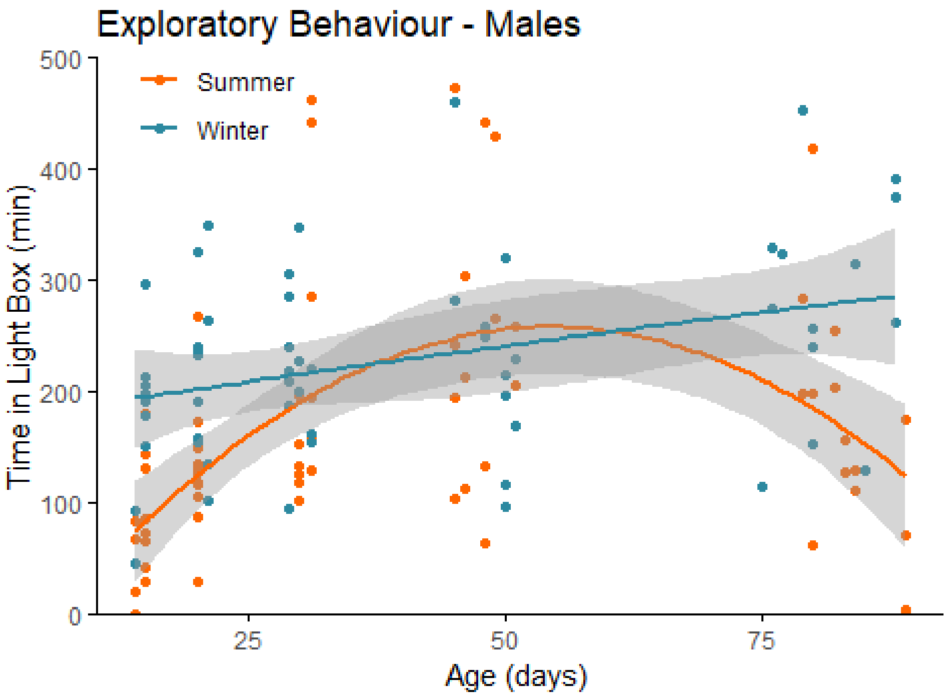


Effect of season: F=3.55, p=0.00056

This panel illustrates the results of a light-dark box behavioural experiment with male summer and winter hamsters between the ages of 15 and 90 days old. The time spent in the light compartment of the box was quantified as our measure of exploratory behaviour. Summer hamsters are housed under a summer-mimicking long photoperiod, while winter hamsters are housed under a winter-mimicking short photoperiod.

*Summary Statistics:*

| Photo | N | LigDar | sd | se | ci |
|---|---|---|---|---|---|
| Summer | 66 | 167.5938 | 112.06119 | 13.79378 | 27.54808 |
| Winter | 57 | 228.5547 | 88.90156 | 11.77530 | 23.58876 |

We found that male summer hamsters show a different developmental trajectory of exploratory behaviour compared to male winter hamsters. We calculated a polynomial regression with age as the continuous variable, season as a categorical variable, and time in the light box as the response variable.

```
##
## Call:
## lm(formula = LigDar ~ poly(Age.Test, 4) * Photo, data = data[data$Age.Test
<
##     100, ])
```

```
##
## Residuals:
##      Min       1Q   Median       3Q      Max
## -178.032  -52.781   -6.971   36.361  263.231
##
## Coefficients:
##                                Estimate Std. Error t value Pr(>|t|)
## (Intercept)                      169.26      11.25  15.049  < 2e-16 ***
## poly(Age.Test, 4)1               296.03     125.19   2.365  0.01976 *
## poly(Age.Test, 4)2              -595.03     121.15  -4.912 3.07e-06 ***
## poly(Age.Test, 4)3               -64.73     132.16  -0.490  0.62524
## poly(Age.Test, 4)4              -181.46     125.95  -1.441  0.15242
## PhotoWinter                       58.76      16.55   3.551  0.00056 ***
## poly(Age.Test, 4)1:PhotoWinter    69.53     184.85   0.376  0.70753
## poly(Age.Test, 4)2:PhotoWinter   577.15     185.39   3.113  0.00235 **
## poly(Age.Test, 4)3:PhotoWinter   234.84     184.68   1.272  0.20613
## poly(Age.Test, 4)4:PhotoWinter   156.58     184.18   0.850  0.39703
## ---
## Signif. codes:  0 '***' 0.001 '**' 0.01 '*' 0.05 '.' 0.1 ' ' 1
##
## Residual standard error: 91.02 on 113 degrees of freedom
##   (1 observation deleted due to missingness)
## Multiple R-squared:  0.3179, Adjusted R-squared:  0.2636
## F-statistic: 5.851 on 9 and 113 DF,  p-value: 1.134e-06
```

#### Figure 3, panel L

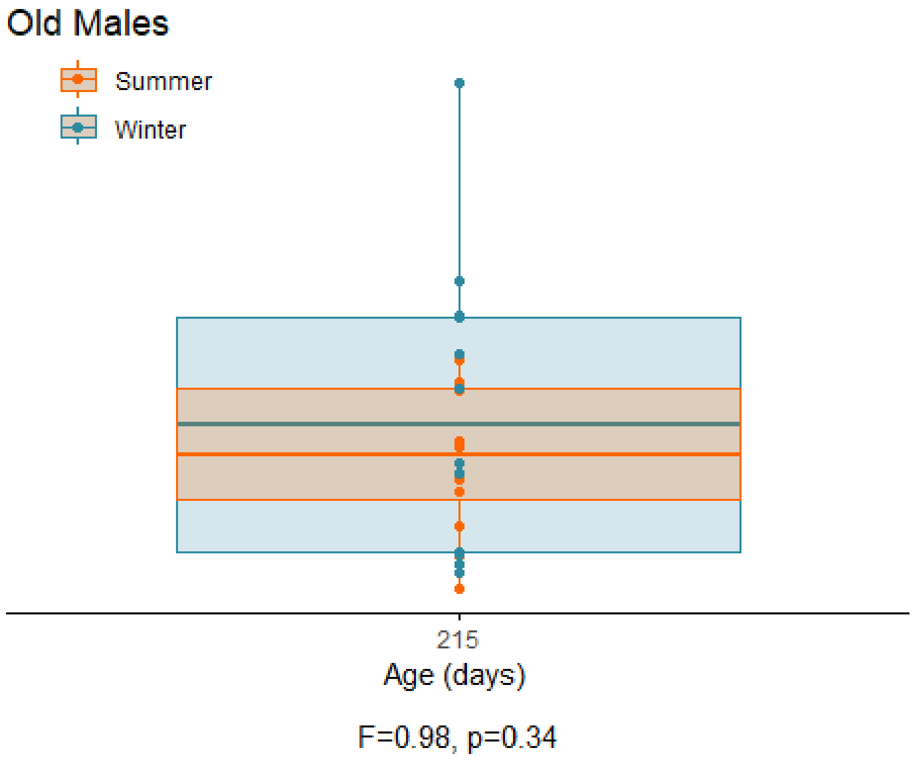


This panel illustrates the results of a light-dark box behavioural experiment with 215-day-old male summer and winter hamsters. The time spent in the light compartment of the box was quantified as our measure of exploratory behaviour. Summer hamsters are housed under a summer-mimicking long photoperiod, while winter hamsters are housed under a winter-mimicking short photoperiod.

*Summary Statistics:*

| Photo | N | LigDar | sd | se | ci |
|---|---|---|---|---|---|
| Summer | 12 | 132.9385 | 69.18096 | 19.97082 | 43.95549 |
| Winter | 12 | 175.8750 | 136.00015 | 39.25986 | 86.41037 |

We found that 215-day-old male summer and winter hamsters show similar levels of exploratory behaviour. We calculated a t-test with season as the independent variable, and time in the light box as the dependent variable.

```
##
## Call:
## lm(formula = LigDar ~ Photo, data = data[data$Age.Test > 100,
##     ])
##
## Residuals:
##     Min      1Q  Median      3Q     Max
## -148.85  -69.75    2.11   71.53  299.86
```

```
##
## Coefficients:
##             Estimate Std. Error t value Pr(>|t|)
## (Intercept)   132.94      31.15   4.268 0.000313 ***
## PhotoWinter    42.94      44.05   0.975 0.340267
## ---
## Signif. codes:  0 '***' 0.001 '**' 0.01 '*' 0.05 '.' 0.1 ' ' 1
##
## Residual standard error: 107.9 on 22 degrees of freedom
##   (8 observations deleted due to missingness)
## Multiple R-squared:  0.0414, Adjusted R-squared:  -0.00217
## F-statistic: 0.9502 on 1 and 22 DF,  p-value: 0.3403
```

#### Figure 4, panel C

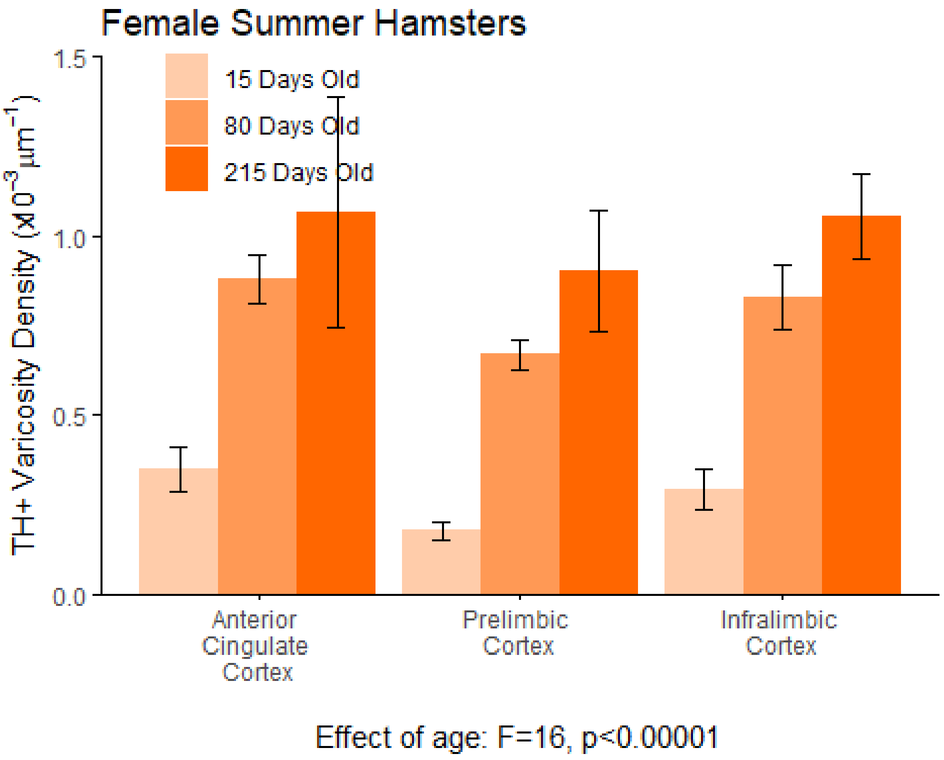


This panel illustrates the results of the quantification of the density of TH-positive varicosities in the dopamine-innervated part of the medial prefrontal cortex. This analysis was conducted in 15-, 80-, and 215-day-old female hamsters that were housed under a summer-mimicking light cycle. The density of dopamine varicosities in each area was quantified using standard unbiased stereology.

*Summary Statistics:*

| age | region | N | density.graph | sd | se | ci |
|---|---|---|---|---|---|---|
| 15 Days Old | Anterior Cingulate Cortex | 6 | 0.35 | 0.15 | 0.06 | 0.16 |
| 15 Days Old | Infralimbic Cortex | 6 | 0.29 | 0.14 | 0.06 | 0.15 |
| 15 Days Old | Prelimbic Cortex | 6 | 0.18 | 0.06 | 0.02 | 0.06 |
| 215 Days Old | Anterior Cingulate Cortex | 4 | 1.07 | 0.65 | 0.32 | 1.03 |
| 215 Days Old | Infralimbic Cortex | 4 | 1.05 | 0.23 | 0.12 | 0.37 |
| 215 Days Old | Prelimbic Cortex | 4 | 0.90 | 0.34 | 0.17 | 0.54 |
| 80 Days Old | Anterior Cingulate Cortex | 8 | 0.88 | 0.20 | 0.07 | 0.16 |
| 80 Days Old | Infralimbic Cortex | 8 | 0.83 | 0.26 | 0.09 | 0.21 |
| 80 Days Old | Prelimbic Cortex | 8 | 0.67 | 0.12 | 0.04 | 0.10 |

Using an analysis of variance (ANOVA), we found that the density of dopamine varicosities in the medial prefrontal cortex increases with age in summer female hamsters. We calculated a mixed-effects ANOVA with brain region, age, and the time each sample was stored in the fridge before staining as fixed effects, hamster ID as a random effect, and varicosity density as the response variable.

```
## 
## Error: hamster_ID
##     Df Sum Sq Mean Sq
## age  1 0.3055  0.3055
## 
## Error: hamster_ID:region
##        Df Sum Sq Mean Sq
## region  2  2.911   1.456
## 
## Error: Within
##                              Df Sum Sq Mean Sq F value   Pr(>F)    
## region                        2   0.17   0.085   0.128 0.879912    
## age                           2  22.27  11.135  16.719  1.5e-06 ***
## fridge.scaled.age             1  10.97  10.970  16.472 0.000139 ***
## region:age                    4   0.90   0.224   0.337 0.852090    
## region:fridge.scaled.age      2   0.44   0.222   0.333 0.717797    
## age:fridge.scaled.age         2   2.73   1.365   2.049 0.137353    
## region:age:fridge.scaled.age  4   0.35   0.086   0.130 0.971087    
## Residuals                    63  41.96   0.666                     
## ---
## Signif. codes:  0 '***' 0.001 '**' 0.01 '*' 0.05 '.' 0.1 ' ' 1
```

#### Figure 4, panel D

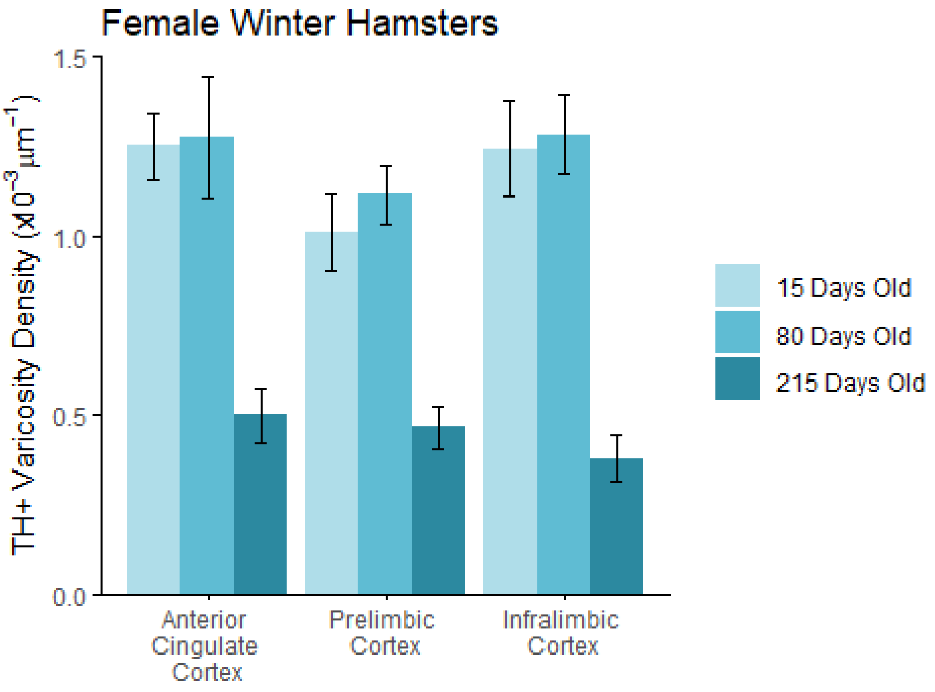


Effect of age: F=12, p=0.00004

This panel illustrates the results of the quantification of the density of TH-positive varicosities in the dopamine-innervated part of the medial prefrontal cortex. This analysis was conducted in 15-, 80-, and 215-day-old female hamsters that were housed under a winter-mimicking light cycle. The density of dopamine varicosities in each area was quantified using standard unbiased stereology.

*Summary Statistics:*

| age | region | N | density.graph | sd | se | ci |
|---|---|---|---|---|---|---|
| 15 Days Old | Anterior Cingulate Cortex | 8 | 1.25 | 0.27 | 0.10 | 0.23 |
| 15 Days Old | Infralimbic Cortex | 8 | 1.24 | 0.37 | 0.13 | 0.31 |
| 15 Days Old | Prelimbic Cortex | 8 | 1.01 | 0.30 | 0.11 | 0.25 |
| 215 Days Old | Anterior Cingulate Cortex | 8 | 0.50 | 0.22 | 0.08 | 0.18 |
| 215 Days Old | Infralimbic Cortex | 8 | 0.38 | 0.18 | 0.06 | 0.15 |
| 215 Days Old | Prelimbic Cortex | 8 | 0.46 | 0.16 | 0.06 | 0.14 |
| 80 Days Old | Anterior Cingulate Cortex | 8 | 1.28 | 0.48 | 0.17 | 0.40 |
| 80 Days Old | Infralimbic Cortex | 8 | 1.28 | 0.31 | 0.11 | 0.26 |
| 80 Days Old | Prelimbic Cortex | 8 | 1.11 | 0.23 | 0.08 | 0.19 |

Using an analysis of variance (ANOVA), we found that the density of dopamine varicosities in the medial prefrontal cortex decreases with age in winter female hamsters. We calculated a mixed-effects ANOVA with brain region, age, and the time each sample was stored in the fridge before staining as fixed effects, hamster ID as a random effect, and varicosity density as the response variable.

```
## 
## Error: hamster_ID
##     Df Sum Sq Mean Sq
## age  1  0.224   0.224
## 
## Error: hamster_ID:region
##        Df Sum Sq Mean Sq
## region  2 0.1102 0.05511
## 
## Error: Within
##                             Df Sum Sq Mean Sq F value   Pr(>F)    
## region                       2  0.014  0.0068   0.056 0.945370    
## age                          2  2.969  1.4845  12.331 4.28e-05 ***
## fridge.scaled.age            1  0.027  0.0265   0.220 0.640773    
## region:age                   4  0.044  0.0109   0.091 0.984992    
## region:fridge.scaled.age     2  0.005  0.0025   0.021 0.979068    
## age:fridge.scaled.age        2  2.405  1.2027   9.991 0.000218 ***
## region:age:fridge.scaled.age  4  0.075  0.0187   0.155 0.959819    
## Residuals                   51  6.139  0.1204                     
## ---
## Signif. codes:  0 '***' 0.001 '**' 0.01 '*' 0.05 '.' 0.1 ' ' 1
```

#### Figure 4, panel E

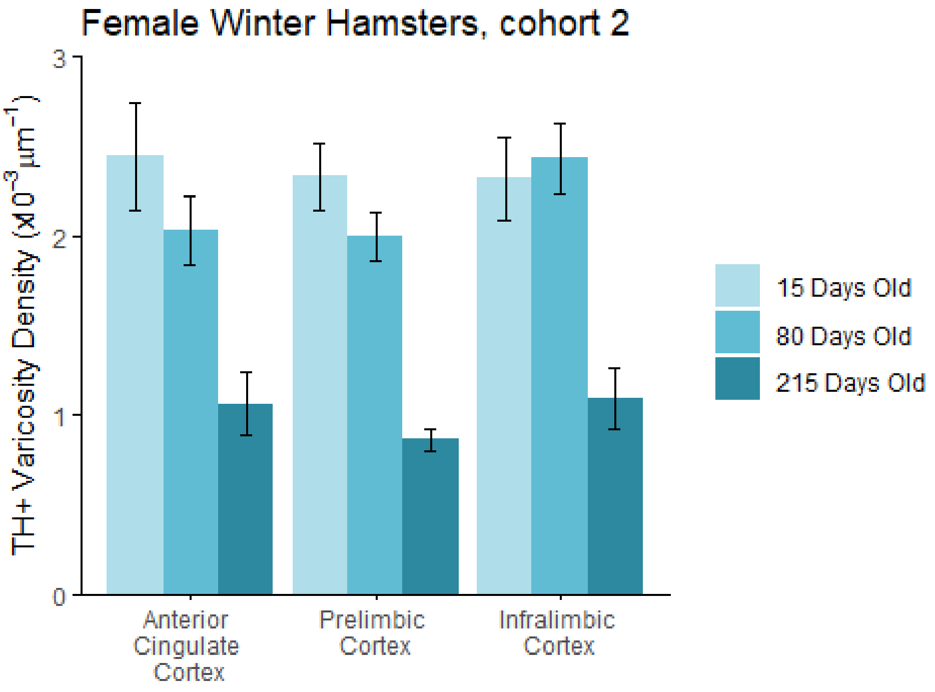


Effect of age: F=35, p<0.00001

This panel illustrates the results of the quantification of the density of TH-positive varicosities in the dopamine-innervated part of the medial prefrontal cortex. This analysis was conducted in 15-, 80-, and 215-day-old female hamsters that were housed under a winter-mimicking light cycle. The density of dopamine varicosities in each area was quantified using standard unbiased stereology. This is an independent cohort of hamsters from those represented in the preceding panel.

*Summary Statistics:*

| age | region | N | density.graph | sd | se | ci |
|---|---|---|---|---|---|---|
| 15 Days Old | Anterior Cingulate Cortex | 8 | 2.44 | 0.85 | 0.30 | 0.71 |
| 15 Days Old | Infralimbic Cortex | 8 | 2.32 | 0.66 | 0.23 | 0.55 |
| 15 Days Old | Prelimbic Cortex | 8 | 2.33 | 0.53 | 0.19 | 0.44 |
| 215 Days Old | Anterior Cingulate Cortex | 8 | 1.06 | 0.50 | 0.18 | 0.42 |
| 215 Days Old | Infralimbic Cortex | 7 | 1.09 | 0.45 | 0.17 | 0.42 |
| 215 Days Old | Prelimbic Cortex | 7 | 0.86 | 0.17 | 0.06 | 0.15 |
| 80 Days Old | Anterior Cingulate Cortex | 8 | 2.03 | 0.54 | 0.19 | 0.45 |
| 80 Days Old | Infralimbic Cortex | 8 | 2.43 | 0.57 | 0.20 | 0.47 |
| 80 Days Old | Prelimbic Cortex | 8 | 1.99 | 0.38 | 0.13 | 0.32 |

Using an analysis of variance (ANOVA), we replicated our finding that the density of dopamine varicosities in the medial prefrontal cortex decreases with age in female winter hamsters. We calculated a mixed-effects ANOVA with brain region, age, and the time each sample was stored in the fridge before staining as fixed effects, hamster ID as a random effect, and varicosity density as the response variable.

```
##
## Error: hamster_ID
##        Df Sum Sq Mean Sq
## region  1 0.3547  0.3547
##
## Error: hamster_ID:region
##        Df Sum Sq Mean Sq
## region  2  0.121 0.06048
##
## Error: Within
##                              Df Sum Sq Mean Sq F value   Pr(>F)
## region                        2  0.115   0.057   0.424    0.657
## age                           2  9.435   4.718  34.871 3.82e-10 ***
## fridge.scaled.age             1  3.683   3.683  27.225 3.65e-06 ***
## region:age                    4  0.188   0.047   0.347    0.845
## region:fridge.scaled.age      2  0.058   0.029   0.216    0.807
## age:fridge.scaled.age         2  4.655   2.328  17.206 2.19e-06 ***
## region:age:fridge.scaled.age  4  0.038   0.009   0.070    0.991
## Residuals                    49  6.629   0.135
## ---
## Signif. codes:  0 '***' 0.001 '**' 0.01 '*' 0.05 '.' 0.1 ' ' 1
```

#### Figure 4, panel F

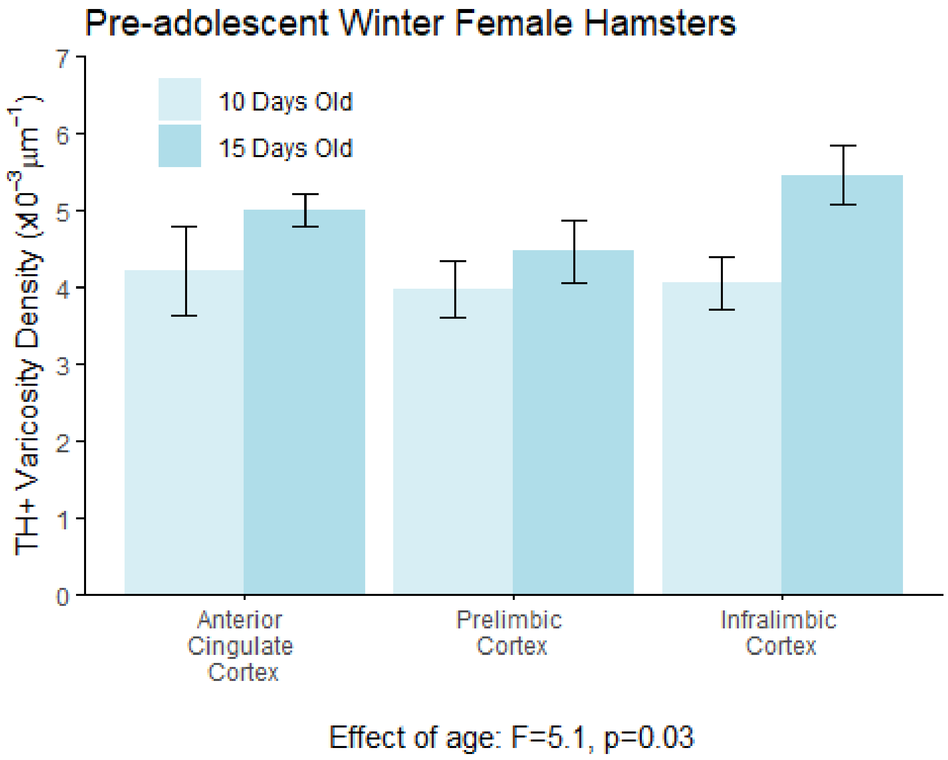


This panel illustrates the results of the quantification of the density of TH-positive varicosities in the dopamine-innervated part of the medial prefrontal cortex. This analysis was conducted in 10- and 15-day-old female hamsters that were housed under a winter-mimicking light cycle. The density of dopamine varicosities in each area was quantified using standard unbiased stereology.

*Summary Statistics:*

| age | region | N | density.graph | sd | se | ci |
|---|---|---|---|---|---|---|
| 10 Days Old | Anterior Cingulate Cortex | 10 | 4.21 | 1.85 | 0.59 | 1.32 |
| 10 Days Old | Infralimbic Cortex | 10 | 4.05 | 1.05 | 0.33 | 0.75 |
| 10 Days Old | Prelimbic Cortex | 10 | 3.97 | 1.17 | 0.37 | 0.84 |
| 15 Days Old | Anterior Cingulate Cortex | 8 | 5.01 | 0.60 | 0.21 | 0.50 |
| 15 Days Old | Infralimbic Cortex | 7 | 5.46 | 1.00 | 0.38 | 0.92 |
| 15 Days Old | Prelimbic Cortex | 7 | 4.46 | 1.07 | 0.41 | 0.99 |

Using an analysis of variance (ANOVA), we found that the density of dopamine varicosities in the medial prefrontal cortex increases between these ages in these winter female hamsters. We calculated a mixed-effects ANOVA with brain region, age, and the time each sample was stored in the fridge before staining as fixed effects, hamster ID as a random effect, and varicosity density as the response variable.

```
##
## Error: hamster_ID
##        Df Sum Sq Mean Sq
## region  1   1.14    1.14
##
## Error: hamster_ID:region
##        Df Sum Sq Mean Sq
## region  2 0.6813  0.3407
##
## Error: Within
##                           Df Sum Sq Mean Sq F value Pr(>F)
## region                     2  0.160  0.0802   0.132 0.8769
## age                        1  3.075  3.0749   5.050 0.0302 *
## fridge.scaled.age          1  0.900  0.9002   1.479 0.2311
## region:age                 2  1.008  0.5041   0.828 0.4443
## region:fridge.scaled.age  2  0.397  0.1987   0.326 0.7235
## Residuals                 40 24.354  0.6089
## ---
## Signif. codes:  0 '***' 0.001 '**' 0.01 '*' 0.05 '.' 0.1 ' ' 1
```

#### Figure 4, panel O

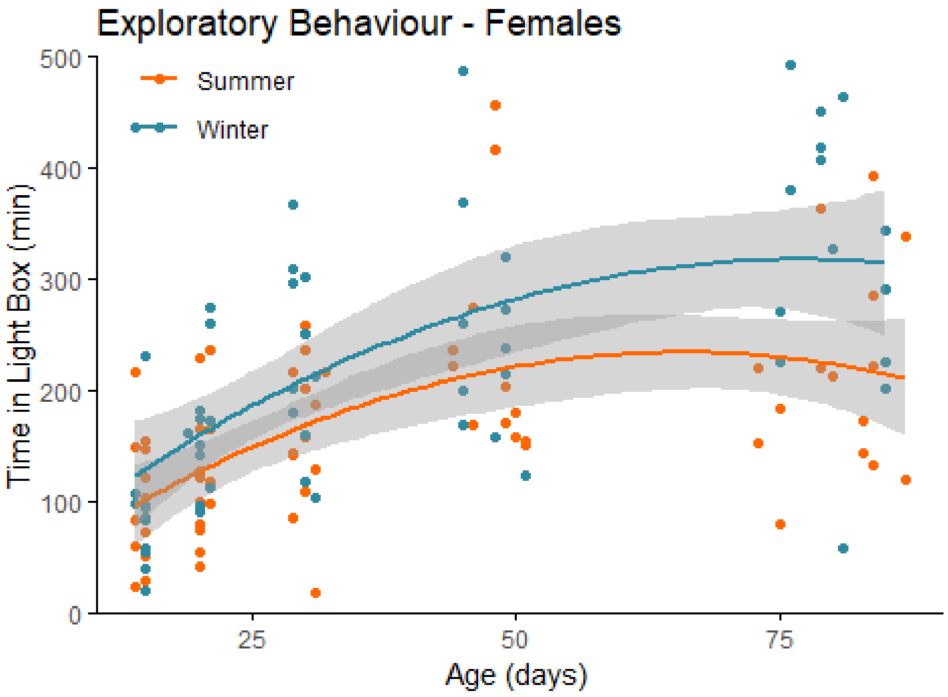


effect of season: F=3.31, p=0.0013

This panel illustrates the results of a light-dark box behavioural experiment with female summer and winter hamsters between the ages of 15 and 90 days old. The time spent in the light compartment of the box was quantified as our measure of exploratory behaviour. Summer hamsters are housed under a summer-mimicking long photoperiod, while winter hamsters are housed under a winter-mimicking short photoperiod.

*Summary Statistics:*

| Photo | N | LigDar | sd | se | ci |
|---|---|---|---|---|---|
| Summer | 66 | 168.8911 | 90.80428 | 11.17723 | 22.32248 |
| Winter | 61 | 221.6236 | 121.24855 | 15.52429 | 31.05320 |

We found that female summer hamsters show a similar developmental trajectory of exploratory behaviour compared to female winter hamsters across adolescence, however the magnitude of the behavioural shifts were larger in the winter hamsters. We calculated a polynomial regression with age as the continuous variable, season as a categorical variable, and time in the light box as the response variable.

```
##
## Call:
## lm(formula = LigDar ~ poly(Age.Test, 4) * Photo, data = data[data$Age.Test
<
##     100, ])
##
```

```
## Residuals:
##     Min      1Q  Median      3Q     Max
## -267.70  -56.93  -10.31   47.97  332.79
##
## Coefficients:
##                                Estimate Std. Error t value Pr(>|t|)
## (Intercept)                     168.396     10.915  15.428  < 2e-16 ***
## poly(Age.Test, 4)1              465.122    123.087   3.779 0.000249 ***
## poly(Age.Test, 4)2             -267.612    122.129  -2.191 0.030417 *
## poly(Age.Test, 4)3               70.392    117.973   0.597 0.551876
## poly(Age.Test, 4)4              140.279    120.792   1.161 0.247874
## PhotoWinter                      52.078     15.757   3.305 0.001260 **
## poly(Age.Test, 4)1:PhotoWinter  269.615    178.563   1.510 0.133761
## poly(Age.Test, 4)2:PhotoWinter    1.191    178.557   0.007 0.994688
## poly(Age.Test, 4)3:PhotoWinter  -86.713    179.930  -0.482 0.630759
## poly(Age.Test, 4)4:PhotoWinter -384.278    179.857  -2.137 0.034716 *
## ---
## Signif. codes:  0 '***' 0.001 '**' 0.01 '*' 0.05 '.' 0.1 ' ' 1
##
## Residual standard error: 88.54 on 117 degrees of freedom
## Multiple R-squared:  0.391,  Adjusted R-squared:  0.3442
## F-statistic: 8.348 on 9 and 117 DF,  p-value: 1.542e-09
```

#### Figure 4, panel P

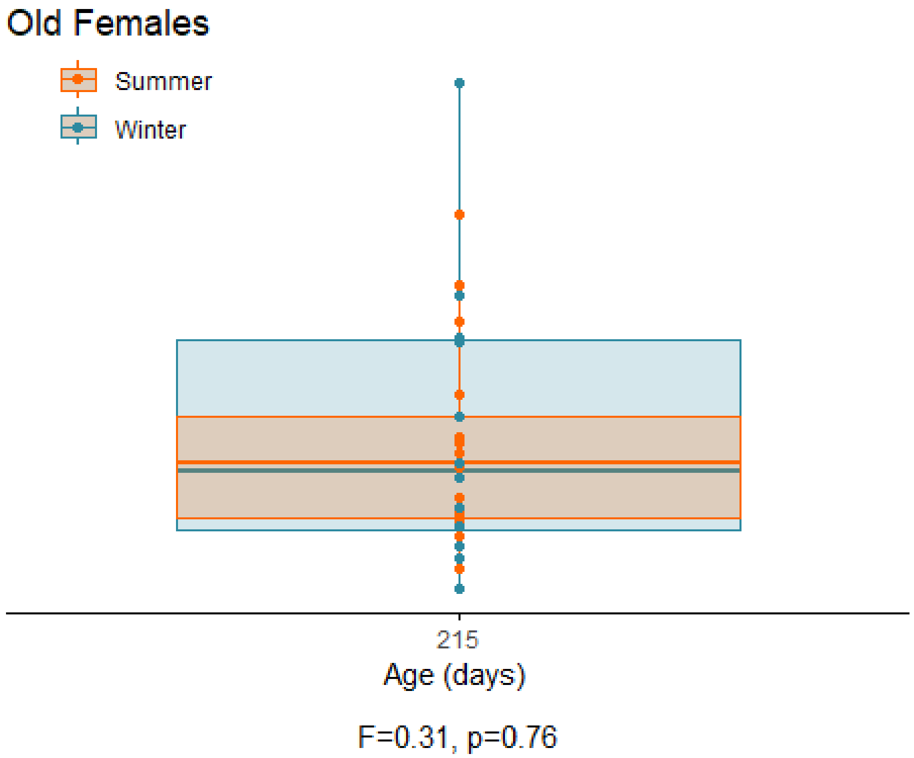


This panel illustrates the results of a light-dark box behavioural experiment with 215-day-old female summer and winter hamsters. The time spent in the light compartment of the box was quantified as our measure of exploratory behaviour. Summer hamsters are housed under a summer-mimicking long photoperiod, while winter hamsters are housed under a winter-mimicking short photoperiod.

*Summary Statistics:*

| Photo | N | LigDar | sd | se | ci |
|---|---|---|---|---|---|
| Summer | 15 | 197.0663 | 91.29994 | 23.57354 | 50.56022 |
| Winter | 12 | 210.3282 | 131.63499 | 37.99975 | 83.63688 |

We found that 215-day-old female summer and winter hamsters show similar levels of exploratory behaviour. We calculated a t-test with season as the independent variable, and time in the light box as the dependent variable.

```
##
## Call:
## lm(formula = LigDar ~ Photo, data = data[data$Age.Test > 100,
##     ])
##
## Residuals:
##     Min      1Q  Median      3Q     Max
## -145.47  -73.59  -22.96   61.43  315.42
```

```
##
## Coefficients:
##             Estimate Std. Error t value Pr(>|t|)
## (Intercept)   197.07      28.63   6.884 3.24e-07 ***
## PhotoWinter    13.26      42.94   0.309     0.76
## ---
## Signif. codes:  0 '***' 0.001 '**' 0.01 '*' 0.05 '.' 0.1 ' ' 1
##
## Residual standard error: 110.9 on 25 degrees of freedom
##   (8 observations deleted due to missingness)
## Multiple R-squared:  0.003801,   Adjusted R-squared:  -0.03605
## F-statistic: 0.09539 on 1 and 25 DF,  p-value: 0.76
```
