## Supplementary Figures for "The scheduling of adolescence with Netrin-1 and UNC5C"

Daniel Hoops

### Supplementary Figure 1

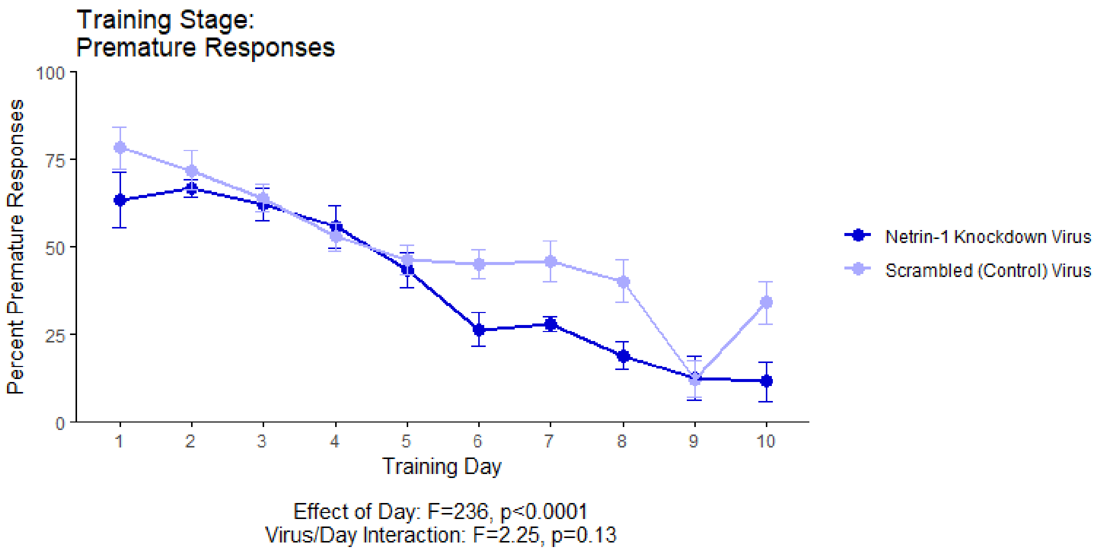


This graph illustrates the ability of adult (60+ day old) mice to perform a learned response to a stimulus in order to receive a food reward as part of the learning phase leading up to the GoNoGo behavioural test. "Percent Premature Responses" quantifies the percentage of trials in which the mice performed the learned response before before the onset of the cue, and therefore ended the trial without receiving a reward. In this experiment, at the onset of adolescence (21 days old) the mice received an injection of a virus that either knocked down Netrin-1 or served as a nonfunctional control. The viruses were injected along the route by which dopamine axons grow from the nucleus accumbens to the medial prefrontal cortex during adolescence. The primary results of this experiment are presented in the main manuscript as figure panels 1M-1Q.

*Summary Statistics:*

| virus | Day | N | Percent.Premature | sd | se | ci |
|---|---|---|---|---|---|---|
| Netrin-1 Knockdown Virus | 1 | 10 | 0.63 | 0.25 | 0.08 | 0.18 |
| Netrin-1 Knockdown Virus | 2 | 10 | 0.67 | 0.08 | 0.03 | 0.06 |
| Netrin-1 Knockdown Virus | 3 | 10 | 0.62 | 0.14 | 0.05 | 0.10 |
| Netrin-1 Knockdown Virus | 4 | 10 | 0.56 | 0.19 | 0.06 | 0.14 |
| Netrin-1 Knockdown Virus | 5 | 10 | 0.43 | 0.16 | 0.05 | 0.11 |
| Netrin-1 Knockdown Virus | 6 | 10 | 0.26 | 0.15 | 0.05 | 0.11 |
| Netrin-1 Knockdown Virus | 7 | 10 | 0.28 | 0.06 | 0.02 | 0.05 |

| virus | Day | N | Percent.Premature | sd | se | ci |
|---|---|---|---|---|---|---|
| Netrin-1 Knockdown Virus | 8 | 10 | 0.19 | 0.13 | 0.04 | 0.09 |
| Netrin-1 Knockdown Virus | 9 | 10 | 0.13 | 0.19 | 0.06 | 0.14 |
| Netrin-1 Knockdown Virus | 10 | 10 | 0.12 | 0.17 | 0.06 | 0.12 |
| Scrambled (Control) Virus | 1 | 10 | 0.78 | 0.19 | 0.06 | 0.14 |
| Scrambled (Control) Virus | 2 | 10 | 0.72 | 0.18 | 0.06 | 0.13 |
| Scrambled (Control) Virus | 3 | 10 | 0.64 | 0.13 | 0.04 | 0.09 |
| Scrambled (Control) Virus | 4 | 10 | 0.53 | 0.13 | 0.04 | 0.09 |
| Scrambled (Control) Virus | 5 | 10 | 0.46 | 0.13 | 0.04 | 0.09 |
| Scrambled (Control) Virus | 6 | 10 | 0.45 | 0.13 | 0.04 | 0.09 |
| Scrambled (Control) Virus | 7 | 10 | 0.46 | 0.18 | 0.06 | 0.13 |
| Scrambled (Control) Virus | 8 | 10 | 0.40 | 0.19 | 0.06 | 0.14 |
| Scrambled (Control) Virus | 9 | 10 | 0.12 | 0.16 | 0.05 | 0.12 |
| Scrambled (Control) Virus | 10 | 10 | 0.34 | 0.19 | 0.06 | 0.14 |

An analysis of variance (ANOVA) revealed no significant differences between viral treatments in the percent of responses that were premature. We determined this using a mixed-effects ANOVA with virus and day as fixed effects, mouse ID as a random effect, and percent of premature responses as the response variable.

```
## Analysis of Deviance Table (Type II tests)
##
## Response: Percent.Premature
##              Chisq Df Pr(>Chisq)
## Day       235.7508  1     <2e-16 ***
## virus       2.1679  1     0.1409
## Day:virus   2.2534  1     0.1333
## ---
## Signif. codes:  0 '***' 0.001 '**' 0.01 '*' 0.05 '.' 0.1 ' ' 1
```

### Supplementary Figure 2

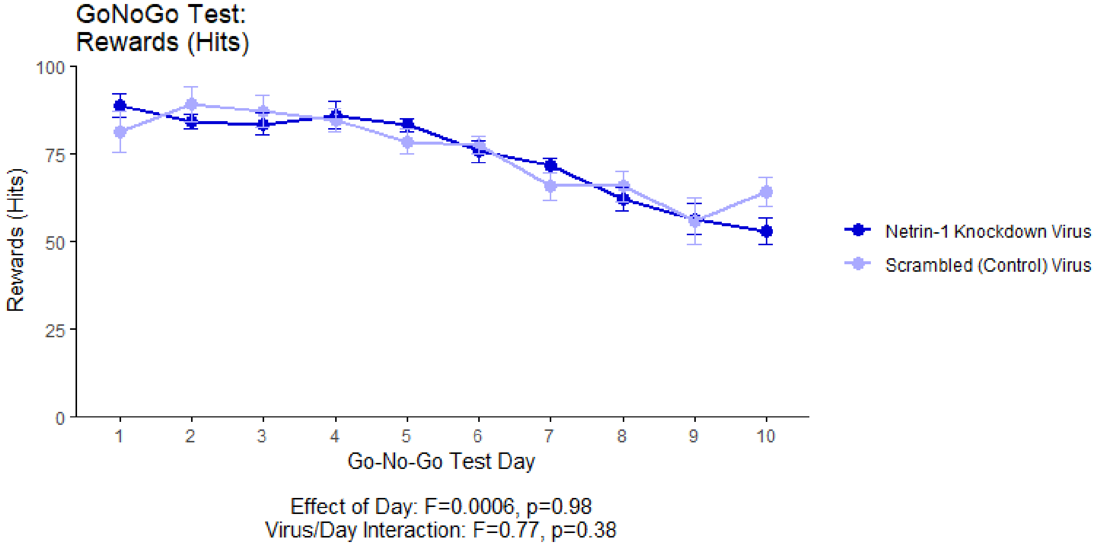


This graph illustrates the ability of adult (60+ day old) mice to perform a learned response to a stimulus in order to receive a food reward as part of the GoNoGo behavioural test. "Hits" quantifies the percentage of trials in which the mice performed correctly in response to a visual cue and in the absence of an auditory cue (a "hit"), and received a food reward as a result.

In this experiment, at the onset of adolescence (21 days old) the mice received an injection of a virus that either knocked down Netrin-1 or served as a nonfunctional control. The viruses were injected along the route by which dopamine axons grow from the nucleus accumbens to the mPFC during adolescence. The primary results of this experiment are presented in the main manuscript as figure panels 1M-1Q.

*Summary Statistics:*

| virus | Day | N | Hits | sd | se | ci |
|---|---|---|---|---|---|---|
| Netrin-1 Knockdown Virus | 1 | 10 | 0.89 | 0.11 | 0.03 | 0.08 |
| Netrin-1 Knockdown Virus | 2 | 10 | 0.84 | 0.07 | 0.02 | 0.05 |
| Netrin-1 Knockdown Virus | 3 | 10 | 0.84 | 0.10 | 0.03 | 0.07 |
| Netrin-1 Knockdown Virus | 4 | 10 | 0.86 | 0.13 | 0.04 | 0.09 |
| Netrin-1 Knockdown Virus | 5 | 10 | 0.83 | 0.06 | 0.02 | 0.05 |
| Netrin-1 Knockdown Virus | 6 | 10 | 0.76 | 0.10 | 0.03 | 0.07 |
| Netrin-1 Knockdown Virus | 7 | 10 | 0.72 | 0.07 | 0.02 | 0.05 |
| Netrin-1 Knockdown Virus | 8 | 10 | 0.62 | 0.11 | 0.03 | 0.08 |
| Netrin-1 Knockdown Virus | 9 | 10 | 0.56 | 0.14 | 0.04 | 0.10 |
| Netrin-1 Knockdown Virus | 10 | 8 | 0.53 | 0.11 | 0.04 | 0.09 |
| Scrambled (Control) Virus | 1 | 10 | 0.81 | 0.18 | 0.06 | 0.13 |

| virus | Day | N | Hits | sd | se | ci |
|---|---|---|---|---|---|---|
| Scrambled (Control) Virus | 2 | 10 | 0.89 | 0.16 | 0.05 | 0.11 |
| Scrambled (Control) Virus | 3 | 10 | 0.87 | 0.14 | 0.05 | 0.10 |
| Scrambled (Control) Virus | 4 | 10 | 0.85 | 0.10 | 0.03 | 0.07 |
| Scrambled (Control) Virus | 5 | 10 | 0.79 | 0.12 | 0.04 | 0.08 |
| Scrambled (Control) Virus | 6 | 10 | 0.78 | 0.08 | 0.02 | 0.06 |
| Scrambled (Control) Virus | 7 | 10 | 0.66 | 0.13 | 0.04 | 0.09 |
| Scrambled (Control) Virus | 8 | 10 | 0.66 | 0.14 | 0.04 | 0.10 |
| Scrambled (Control) Virus | 9 | 10 | 0.56 | 0.21 | 0.07 | 0.15 |
| Scrambled (Control) Virus | 10 | 10 | 0.64 | 0.13 | 0.04 | 0.09 |

An analysis of variance (ANOVA) revealed no significant differences between viral treatments in the ability of the mice to respond correctly to a visual cue. We determined this using a mixed-effects ANOVA with virus and day as fixed effects, mouse ID as a random effect, and percent of responses that were hits as the response variable.

```
## Analysis of Deviance Table (Type II tests)
##
## Response: Percent.Omission.Errors
##              Chisq Df Pr(>Chisq)
## Day       128.8140  1     <2e-16 ***
## virus       0.0006  1     0.9803
## Day:virus   0.7673  1     0.3811
## ---
## Signif. codes:  0 '***' 0.001 '**' 0.01 '*' 0.05 '.' 0.1 ' ' 1
```

### Supplementary Figure 3

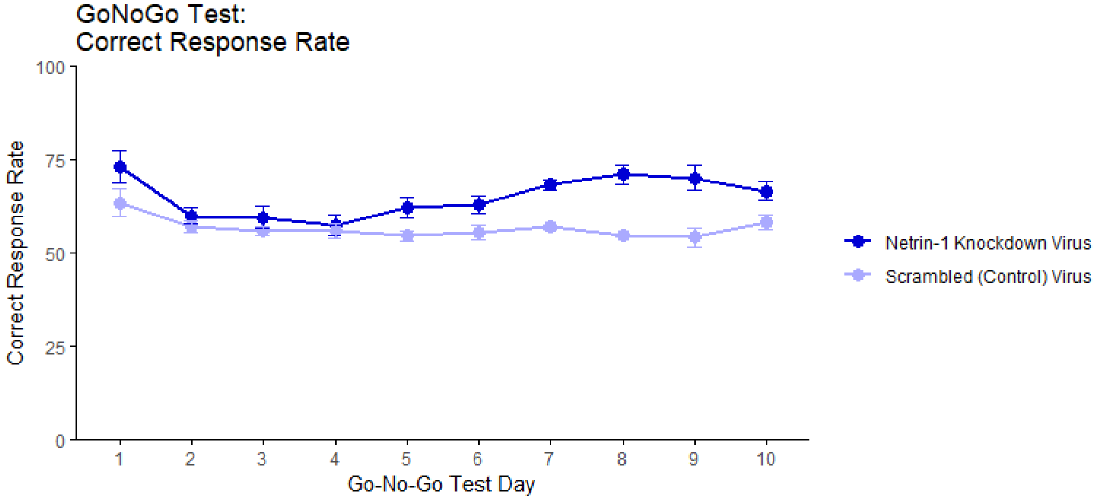


Virus/Day Interaction: F=7.21, p=0.0072

This graph illustrates the ability of adult (60+ day old) mice to perform both "Go" trials and "NoGo" trials correctly. A "Go" trial requires a behavioural response to a visual cue. A "NoGo" trial requires the inhibition of the behavioural response to the visual cue when it is presented with a second, auditory cue. The "Correct Response Rate" quantifies the trials where the mice respond correctly whether the trial is a "Go" trial or a "NoGo" trial (Cuesta et al., 2019). In this experiment, at the onset of adolescence (21 days old) the mice received an injection of a virus that either knocked down Netrin-1 or served as a nonfunctional control. The viruses were injected along the route by which dopamine axons grow from the nucleus accumbens to the medial prefrontal cortex during adolescence. The primary results of this experiment are presented in the main manuscript as figure panels 1M-1Q.

*Summary Statistics:*

| virus | Day | N | Correct.Response.Rate | sd | se | ci |
|---|---|---|---|---|---|---|
| Netrin-1 Knockdown Virus | 1 | 10 | 0.73 | 0.14 | 0.04 | 0.10 |
| Netrin-1 Knockdown Virus | 2 | 10 | 0.60 | 0.07 | 0.02 | 0.05 |
| Netrin-1 Knockdown Virus | 3 | 10 | 0.60 | 0.09 | 0.03 | 0.06 |
| Netrin-1 Knockdown Virus | 4 | 10 | 0.57 | 0.09 | 0.03 | 0.07 |
| Netrin-1 Knockdown Virus | 5 | 10 | 0.62 | 0.08 | 0.03 | 0.06 |
| Netrin-1 Knockdown Virus | 6 | 10 | 0.63 | 0.07 | 0.02 | 0.05 |
| Netrin-1 Knockdown Virus | 7 | 10 | 0.68 | 0.04 | 0.01 | 0.03 |
| Netrin-1 Knockdown Virus | 8 | 10 | 0.71 | 0.09 | 0.03 | 0.06 |
| Netrin-1 Knockdown Virus | 9 | 10 | 0.70 | 0.10 | 0.03 | 0.07 |
| Netrin-1 Knockdown Virus | 10 | 8 | 0.67 | 0.07 | 0.02 | 0.06 |
| Scrambled (Control) Virus | 1 | 10 | 0.64 | 0.12 | 0.04 | 0.09 |
| Scrambled (Control) Virus | 2 | 10 | 0.57 | 0.05 | 0.02 | 0.04 |

| virus | Day | N | Correct.Response.Rate | sd | se | ci |
|---|---|---|---|---|---|---|
| Scrambled (Control) Virus | 3 | 10 | 0.56 | 0.04 | 0.01 | 0.03 |
| Scrambled (Control) Virus | 4 | 10 | 0.56 | 0.06 | 0.02 | 0.04 |
| Scrambled (Control) Virus | 5 | 10 | 0.55 | 0.04 | 0.01 | 0.03 |
| Scrambled (Control) Virus | 6 | 10 | 0.55 | 0.06 | 0.02 | 0.04 |
| Scrambled (Control) Virus | 7 | 10 | 0.57 | 0.02 | 0.01 | 0.02 |
| Scrambled (Control) Virus | 8 | 10 | 0.55 | 0.01 | 0.00 | 0.01 |
| Scrambled (Control) Virus | 9 | 10 | 0.54 | 0.08 | 0.03 | 0.06 |
| Scrambled (Control) Virus | 10 | 10 | 0.58 | 0.06 | 0.02 | 0.04 |

An analysis of variance (ANOVA) revealed a significant difference between viral treatments in the ability of the mice to respond correctly to cues during the GoNoGo test. We determined this using a mixed-effects ANOVA with virus and day as fixed effects, mouse ID as a random effect, and percent of responses that were correct as the response variable.

```
## Analysis of Deviance Table (Type II tests)
##
## Response: Correct.Response.Rate
##            Chisq Df Pr(>Chisq)
## Day       0.2907  1   0.589803
## virus     8.3375  1   0.003883 **
## Day:virus 7.2103  1   0.007249 **
## ---
## Signif. codes:  0 '***' 0.001 '**' 0.01 '*' 0.05 '.' 0.1 ' ' 1
```

### Supplementary Figure 4

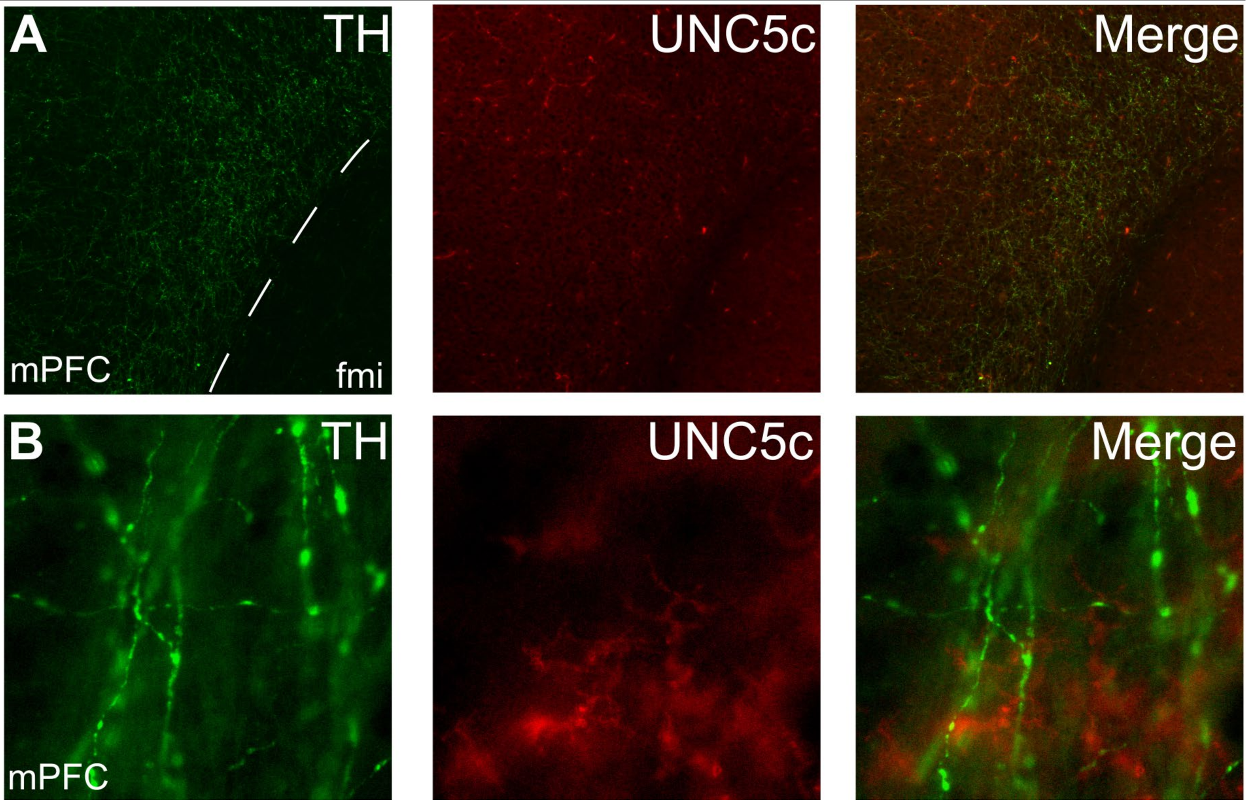


Expression of UNC5c protein in the medial prefrontal cortex of an adult male mouse. Low (A) and high (B) magnification images demonstrate that there is little UNC5c expression in dopamine axons in the medial prefrontal cortex. Here we identify dopamine axons by immunofluorescent staining for tyrosine hydroxylase (TH). Abbreviations: fmi: forceps minor of the corpus callosum, mPFC: medial prefrontal cortex.

### Supplementary Figure 5

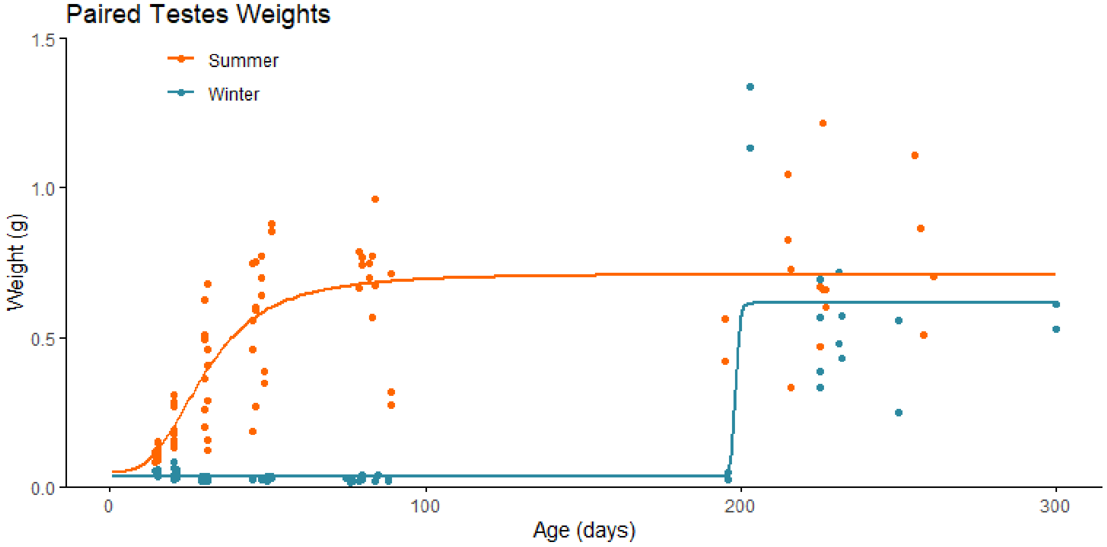


```
##
## Call:
## lm(formula = Gonad.Weight ~ Photo + Age.Test, data = data[!is.na(data[,
##     "Gonad.Weight"]) & data$Sex == "Males", ])
##
## Residuals:
##      Min       1Q   Median       3Q      Max
## -0.44869 -0.13411 -0.02163  0.06710  0.92304
##
## Coefficients:
##               Estimate Std. Error t value Pr(>|t|)
## (Intercept)  0.2897073  0.0274283   10.56   <2e-16 ***
## PhotoWinter -0.3341894  0.0329919  -10.13   <2e-16 ***
## Age.Test     0.0022731  0.0002046   11.11   <2e-16 ***
## ---
## Signif. codes:  0 '***' 0.001 '**' 0.01 '*' 0.05 '.' 0.1 ' ' 1
##
## Residual standard error: 0.2054 on 153 degrees of freedom
## Multiple R-squared:  0.5866, Adjusted R-squared:  0.5812
## F-statistic: 108.6 on 2 and 153 DF,  p-value: < 2.2e-16
```

This graph and linear model illustrate the effects of daylength on reproductive development of male hamsters. Testicular weight is a commonly used proxy for the timing of puberty in male hamsters. The increase in paired testes weight, signaling puberty, is delayed when housed under a winter-mimicking short daylength, compared to a summer-mimicking long daylength. The primary results comparing male summer and winter hamsters are presented in Figure 3.

### Supplementary Figure 6

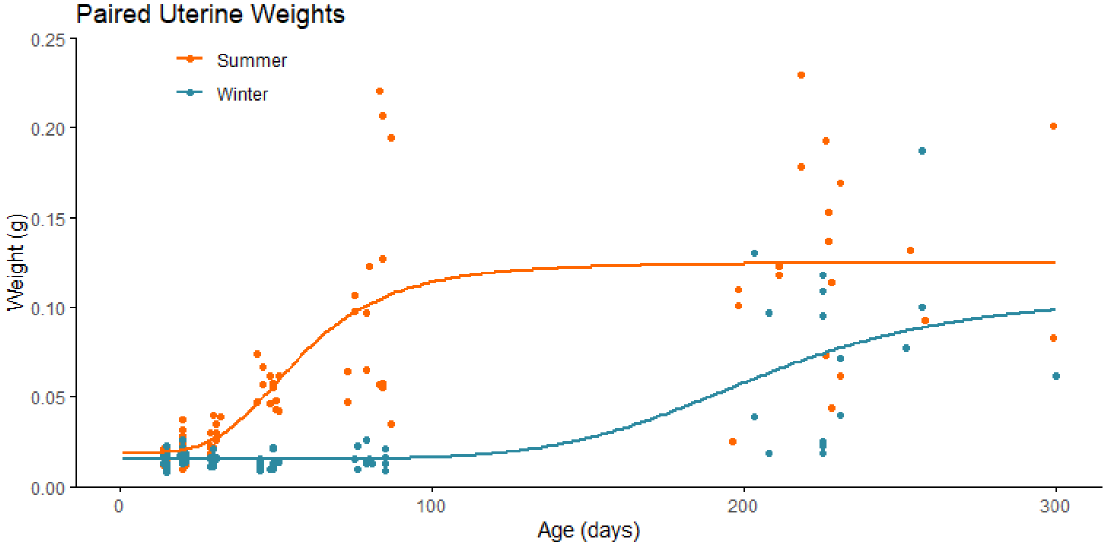


```
##
## Call:
## lm(formula = Gonad.Weight ~ Photo + Age.Test, data = data[!is.na(data[,
##     "Gonad.Weight"]) & data$Sex == "Females", ])
##
## Residuals:
##       Min        1Q    Median        3Q       Max
## -0.082410 -0.017072 -0.003175  0.010316  0.157190
##
## Coefficients:
##               Estimate Std. Error t value Pr(>|t|)
## (Intercept)  3.179e-02  4.595e-03   6.918 1.06e-10 ***
## PhotoWinter -3.484e-02  5.396e-03  -6.456 1.25e-09 ***
## Age.Test     3.858e-04  3.266e-05  11.813  < 2e-16 ***
## ---
## Signif. codes:  0 '***' 0.001 '**' 0.01 '*' 0.05 '.' 0.1 ' ' 1
##
## Residual standard error: 0.0343 on 159 degrees of freedom
## Multiple R-squared:  0.5363, Adjusted R-squared:  0.5305
## F-statistic: 91.96 on 2 and 159 DF,  p-value: < 2.2e-16
```

This graph and linear model illustrate the effects of daylength on the reproductive development of female hamsters. Uterine weight is a commonly used proxy for the timing of puberty in female hamsters. The increase in uterine weight, signalling puberty, is delayed when housed under a winter-mimicking short daylength, compared to a summer-mimicking long daylength. The primary results comparing female summer and winter hamsters are presented in Figure 4.

### Supplementary Figure 7

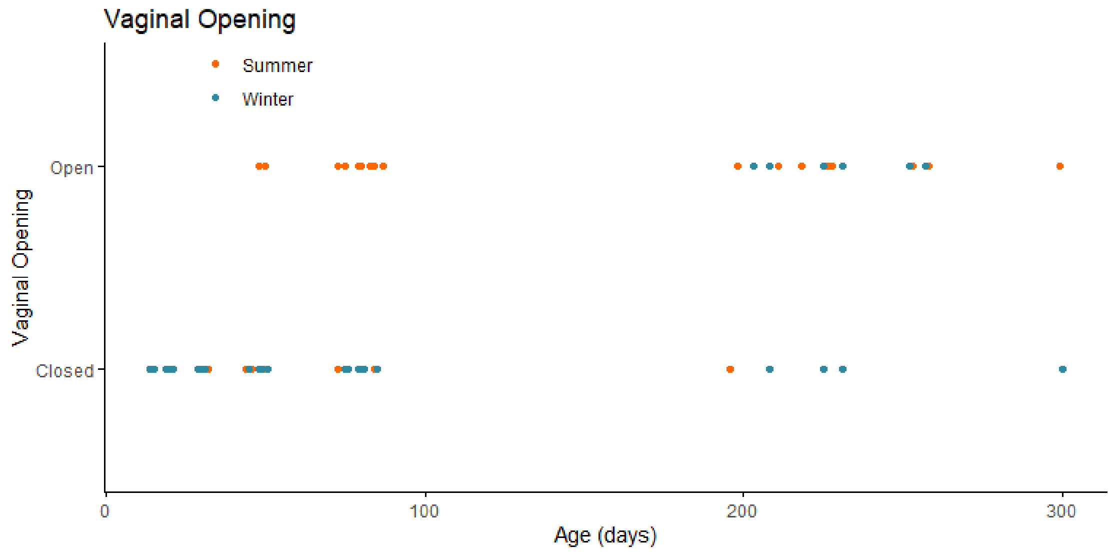


```
##
## Call:
## glm(formula = VaginalOpening ~ Photo + Age.Test, data = data[!is.na(data[,
##     "VaginalOpening"]) & data$Sex == "Females", ])
##
## Deviance Residuals:
##      Min        1Q    Median        3Q       Max
## -0.99156  -0.15022  -0.02265   0.10641   0.74497
##
## Coefficients:
##              Estimate Std. Error t value Pr(>|t|)
## (Intercept)  0.070107   0.038682   1.812   0.0718 .
## PhotoWinter -0.234301   0.045430  -5.157 7.36e-07 ***
## Age.Test     0.003853   0.000275  14.010  < 2e-16 ***
## ---
## Signif. codes:  0 '***' 0.001 '**' 0.01 '*' 0.05 '.' 0.1 ' ' 1
##
## (Dispersion parameter for gaussian family taken to be 0.08335917)
##
##     Null deviance: 32.049  on 161  degrees of freedom
## Residual deviance: 13.254  on 159  degrees of freedom
## AIC: 62.203
##
## Number of Fisher Scoring iterations: 2
```

This graph and generalized linear model illustrate the effects of daylength on the reproductive development of female hamsters. The timing of the opening of the vagina is, in females, a commonly used proxy for the timing of puberty alongside uterine weight. The opening of the vagina, signalling puberty, is delayed when housed under a winter-mimicking short daylength, compared to a summer-mimicking long daylength. The primary results comparing female summer and winter hamsters are presented in Figure 4.

### Supplementary Figure 8

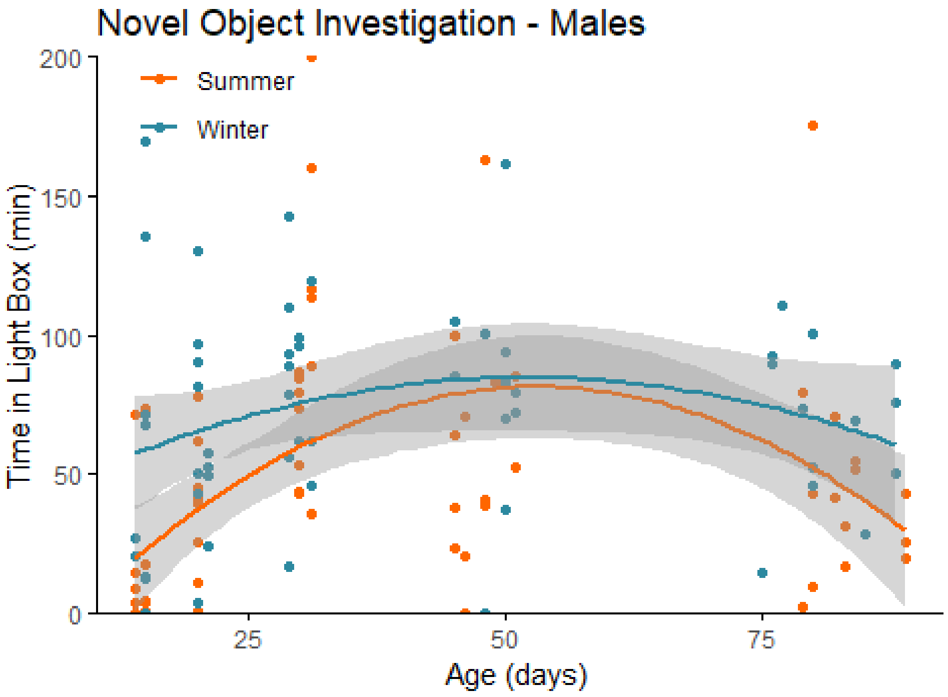


Effect of season: F=2.77, p=0.0065

This panel illustrates the results of a novel object behavioural experiment with male summer and winter hamsters between the ages of 15 and 90 days old. The time spent in the light compartment of the box was quantified as our measure of novel object investigation. Summer hamsters are housed under a summer-mimicking long photoperiod, while winter hamsters are housed under a winter-mimicking short photoperiod.

*Summary Statistics:*

| Photo | N | NovObj | sd | se | ci |
|---|---|---|---|---|---|
| Summer | 66 | 49.42719 | 44.92317 | 5.529660 | 11.04349 |
| Winter | 57 | 70.98883 | 39.29375 | 5.204584 | 10.42603 |

```
##
## Call:
## lm(formula = NovObj ~ poly(Age.Test, 4) * Photo, data = data[data$Age.Test
<
##     100 & data$Sex == "M", ])
##
## Residuals:
##    Min     1Q Median     3Q    Max
## -81.81 -22.71  -2.98  17.17 123.65
##
## Coefficients:
##                                  Estimate Std. Error t value Pr(>|t|)
## (Intercept)                        50.749      4.883  10.393  < 2e-16 ***
```

```
## poly(Age.Test, 4)1                 61.666     53.996   1.142 0.255849
## poly(Age.Test, 4)2               -204.245     52.908  -3.860 0.000189 ***
## poly(Age.Test, 4)3                 99.913     57.107   1.750 0.082907 .
## poly(Age.Test, 4)4                -86.960     54.574  -1.593 0.113856
## PhotoWinter                        19.902      7.179   2.772 0.006513 **
## poly(Age.Test, 4)1:PhotoWinter  -33.556     79.966  -0.420 0.675550
## poly(Age.Test, 4)2:PhotoWinter  104.274     80.642   1.293 0.198635
## poly(Age.Test, 4)3:PhotoWinter  -65.269     79.943  -0.816 0.415961
## poly(Age.Test, 4)4:PhotoWinter   75.718     79.844   0.948 0.344989
## ---
## Signif. codes:  0 '***' 0.001 '**' 0.01 '*' 0.05 '.' 0.1 ' ' 1
##
## Residual standard error: 39.48 on 113 degrees of freedom
##   (1 observation deleted due to missingness)
## Multiple R-squared:  0.2405, Adjusted R-squared:  0.1801
## F-statistic: 3.977 on 9 and 113 DF,  p-value: 0.0001971
```

### Supplementary Figure 9

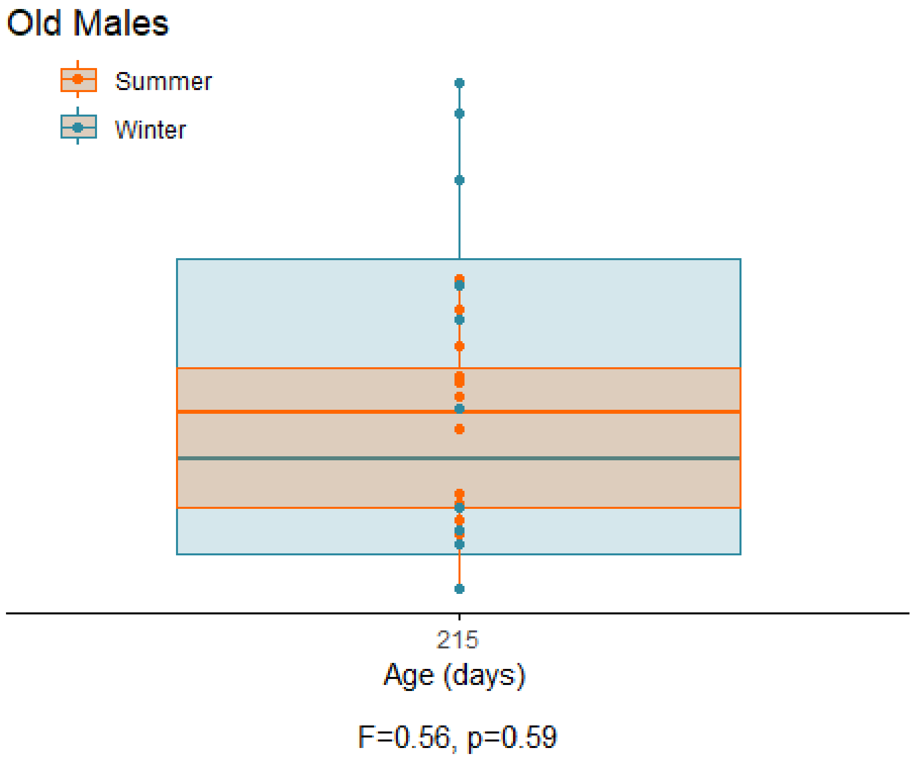


This panel illustrates the results of a novel object behavioural experiment with 215-day-old male summer and winter hamsters. The time spent in the light compartment of the box was quantified as our measure of novel object investigation. Summer hamsters are housed under a summer-mimicking long photoperiod, while winter hamsters are housed under a winter-mimicking short photoperiod.

*Summary Statistics:*

| Photo | N | NovObj | sd | se | ci |
|---|---|---|---|---|---|
| Summer | 12 | 27.21830 | 16.70891 | 4.823448 | 10.61634 |
| Winter | 12 | 33.18527 | 32.99246 | 9.524102 | 20.96241 |

```
##
## Call:
## lm(formula = NovObj ~ Photo, data = data[data$Age.Test > 100 &
##     data$Sex == "M", ])
##
## Residuals:
##    Min     1Q Median     3Q    Max
## -33.19 -20.45  -1.14  15.51  53.73
##
## Coefficients:
##             Estimate Std. Error t value Pr(>|t|)
## (Intercept)   27.218      7.549   3.606  0.00157 **
## PhotoWinter    5.967     10.676   0.559  0.58186
```

```
## ---
## Signif. codes:  0 '***' 0.001 '**' 0.01 '*' 0.05 '.' 0.1 ' ' 1
##
## Residual standard error: 26.15 on 22 degrees of freedom
##   (8 observations deleted due to missingness)
## Multiple R-squared:  0.014,  Adjusted R-squared:  -0.03082
## F-statistic: 0.3124 on 1 and 22 DF,  p-value: 0.5819
```

### Supplementary Figure 10

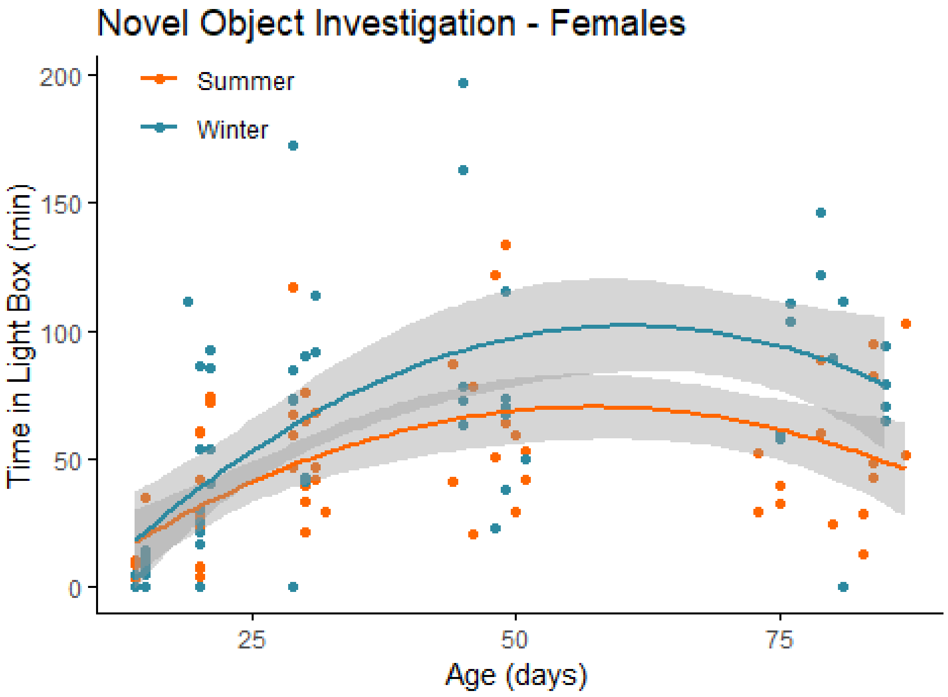


effect of season: F=3.12, p=0.0023

This panel illustrates the results of a novel object behavioural experiment with female summer and winter hamsters between the ages of 15 and 90 days old. The time spent in the light compartment of the box was quantified as our measure of novel object investigation. Summer hamsters are housed under a summer-mimicking long photoperiod, while winter hamsters are housed under a winter-mimicking short photoperiod.

*Summary Statistics:*

| Photo | N | NovObj | sd | se | ci |
|---|---|---|---|---|---|
| Summer | 66 | 44.69326 | 31.92605 | 3.929825 | 7.848403 |
| Winter | 61 | 62.38676 | 47.92103 | 6.135659 | 12.273145 |

```
## 
## Call:
## lm(formula = NovObj ~ poly(Age.Test, 4) * Photo, data = data[data$Age.Test 
< 
##     100 & data$Age.Test < 100 & data$Sex == "F", ])
## 
## Residuals:
##    Min      1Q Median      3Q     Max 
## -90.29 -16.70  -3.14  13.24 114.59 
## 
## Coefficients:
##                                  Estimate Std. Error t value Pr(>|t|)    
## (Intercept)                        44.350      3.982  11.138  < 2e-16 ***
```

```
## poly(Age.Test, 4)1              125.375     44.903   2.792  0.00612 **
## poly(Age.Test, 4)2             -144.834     44.553  -3.251  0.00150 **
## poly(Age.Test, 4)3              108.070     43.037   2.511  0.01340 *
## poly(Age.Test, 4)4                9.230     44.065   0.209  0.83445
## PhotoWinter                      17.958      5.748   3.124  0.00225 **
## poly(Age.Test, 4)1:PhotoWinter  122.753     65.140   1.884  0.06199 .
## poly(Age.Test, 4)2:PhotoWinter  -47.920     65.138  -0.736  0.46340
## poly(Age.Test, 4)3:PhotoWinter  -14.911     65.639  -0.227  0.82069
## poly(Age.Test, 4)4:PhotoWinter -100.221     65.612  -1.527  0.12934
## ---
## Signif. codes:  0 '***' 0.001 '**' 0.01 '*' 0.05 '.' 0.1 ' ' 1
##
## Residual standard error: 32.3 on 117 degrees of freedom
## Multiple R-squared:  0.4295, Adjusted R-squared:  0.3856
## F-statistic: 9.788 on 9 and 117 DF,  p-value: 4.64e-11
```

### Supplementary Figure 11

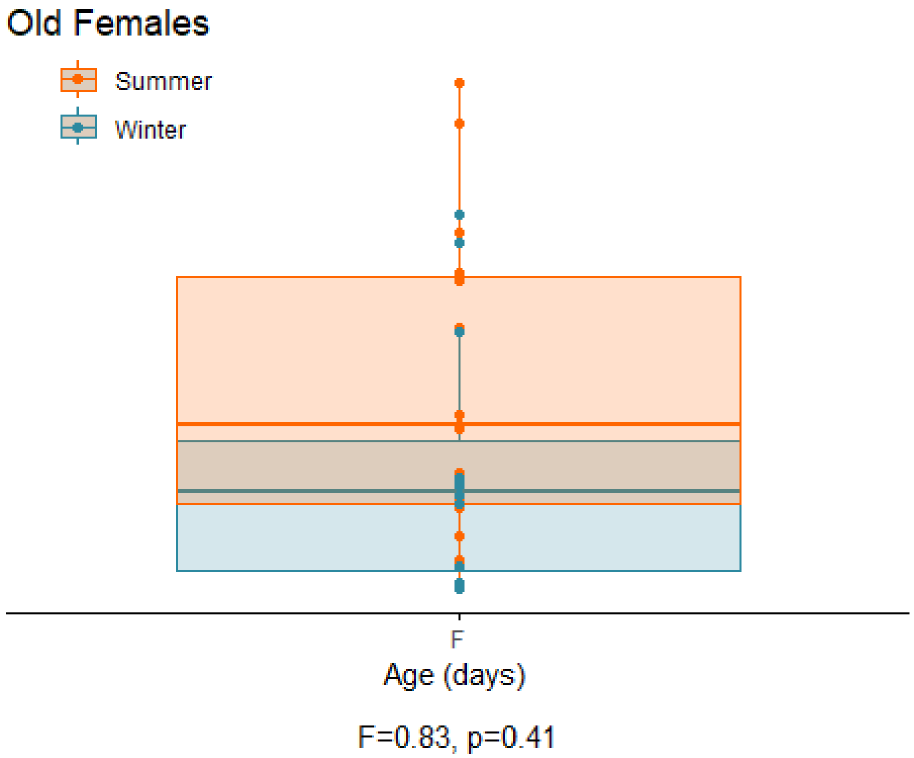


This panel illustrates the results of a novel object behavioural experiment with 215-day-old female summer and winter hamsters. The time spent in the light compartment of the box was quantified as our measure of novel object investigation. Summer hamsters are housed under a summer-mimicking long photoperiod, while winter hamsters are housed under a winter-mimicking short photoperiod.

*Summary Statistics:*

| Photo | N | NovObj | sd | se | ci |
|---|---|---|---|---|---|
| Summer | 27 | 37.53426 | 29.55776 | 5.688393 | 11.69266 |
| Winter | 24 | 30.57994 | 30.59866 | 6.245926 | 12.92068 |

```
##
## Call:
## lm(formula = NovObj ~ Photo, data = data[data$Age.Test > 100,
##     ])
##
## Residuals:
##     Min      1Q  Median      3Q     Max
## -37.534 -24.364  -6.456  18.070  75.810
##
## Coefficients:
##             Estimate Std. Error t value Pr(>|t|)
## (Intercept)   37.534      5.783   6.490 4.09e-08 ***
## PhotoWinter   -6.954      8.431  -0.825    0.413
```

```
## ---
## Signif. codes:  0 '***' 0.001 '**' 0.01 '*' 0.05 '.' 0.1 ' ' 1
##
## Residual standard error: 30.05 on 49 degrees of freedom
##   (16 observations deleted due to missingness)
## Multiple R-squared:  0.0137, Adjusted R-squared:  -0.006432
## F-statistic: 0.6805 on 1 and 49 DF,  p-value: 0.4134
```
